# Time-Resolved Single-Cell Atlas Reveals Early Endothelial Activation and Stage-Dependent Immune–Stromal Communication in HFpEF

**DOI:** 10.64898/2026.08.08.743525

**Authors:** Jiantao Gong, Hannah Morgan, Keara Little, Caleb Cook, Suchandrima Dutta, Riya Bhullar, Olivia Lim, Taliyah Taylor, Rachael Arora, Asma Raja, Yigang Wang, Donald Lynch, Guo-Chang Fan, Wei Huang

## Abstract

**Background:** Heart failure with preserved ejection fraction (HFpEF) is a heterogeneous syndrome associated with metabolic stress, hypertension, systemic inflammation, and microvascular dysfunction. Early cell-type-specific events and intercellular communication programs that accompany disease onset and progression remain poorly defined.

**Methods:** We performed a longitudinal study of HFpEF progression in high-fat diet (HFD)+L-NAME mice at control/baseline (0 weeks, 0w/Ctrl), early (1w), intermediate (4w), and established (8w) stages. Metabolic, hemodynamic, exercise, echocardiographic, and single-cardiomyocyte function were assessed. Cardiac non-cardiomyocytes (non-CMs) were profiled by single-cell RNA sequencing (scRNA-seq), with bulk RNA-seq for tissue-level comparison. Endothelial remodeling was assessed in an L-NAME-independent HFD plus mild transverse aortic constriction model (HFD+mTAC) and a published human HFpEF single-nucleus RNA-seq cohort. An endothelial–macrophage adhesion assay tested whether HFpEF-mimic stress promotes endothelial activation and macrophage adhesion.

**Results:** In the HFD+L-NAME model, metabolic dysfunction, hypertension, reduced exercise tolerance, abnormal diastolic filling with preserved ejection fraction, and altered cardiomyocyte calcium handling were detected by 1w and persisted through 8w. Bulk RNA-seq showed progressive remodeling, with limited change between 8w and 12w, guiding scRNA-seq timepoint selection. scRNA-seq of 94,848 cardiac non-CMs identified nine major populations with stage-dependent remodeling. Endothelial cells (ECs) were recovered in high proportion and showed an early, pronounced transcriptional response, with inflammatory, adhesion, interferon-response, migratory, and vascular-remodeling programs emerging by 1w. Related EC activation signatures were observed in HFD+mTAC and human HFpEF data. Functionally, HFpEF-mimic stress increased adhesion and chemokine expression in human ECs and enhanced macrophage adhesion. Fibroblast matrix remodeling occurred at later stages, while macrophages progressively shifted toward inflammatory states. CellChat suggested stage-dependent communication remodeling from early endothelial–immune interactions toward later macrophage–fibroblast crosstalk.

**Conclusion:** Time-resolved scRNA-seq reveals coordinated, stage-dependent remodeling of the cardiac microvascular and interstitial microenvironment during HFpEF progression. Early endothelial activation emerges before later fibroblast matrix remodeling and inflammatory macrophage remodeling, identifying candidate cell states and signaling pathways for future mechanistic investigation.

**Clinical Perspective:** *What Is New?:*

- This study provides a time-resolved single-cell atlas of the cardiac non-cardiomyocyte compartment across baseline, early, intermediate, and established stages of HFpEF progression, rather than a single late-stage snapshot.
- Endothelial cells exhibit early inflammatory, adhesion, interferon-response, and vascular-remodeling programs within the first week of disease, preceding the later predominance of fibroblast matrix remodeling and inflammatory macrophage remodeling.
- This endothelial activation signature is supported across two mechanistically distinct HFpEF mouse models and aligns with endothelial inflammatory and vascular-remodeling programs in human HFpEF myocardium, supporting its translational relevance.

*What Are the Clinical Implications?:*

- Early endothelial activation may represent a targetable stage of HFpEF pathogenesis that arises before more established structural and fibrotic remodeling.
- Therapeutic strategies aimed at limiting endothelial inflammatory activation or endothelial–immune interactions may help attenuate downstream vascular, immune, and stromal remodeling in HFpEF.
- These findings provide a preclinical foundation for future longitudinal human studies testing whether early endothelial activation can serve as a biomarker, therapeutic target, or disease-staging feature in HFpEF.

## Introduction

Heart failure with preserved ejection fraction (HFpEF) is a major and growing form of heart failure, characterized by impaired ventricular relaxation and exercise intolerance despite preserved systolic function^1–4^. Unlike heart failure with reduced ejection fraction (HFrEF), HFpEF is a multifactorial syndrome associated with hypertension, obesity, metabolic stress, aging, and systemic inflammation. This clinical and biological complexity has made it difficult to identify the early disease-driving mechanisms and to develop effective mechanism-based therapies.

The two-hit mouse model combining high-fat diet (HFD) with L-NAME-mediated nitric oxide synthase inhibition has provided important mechanistic insight into metabolic stress, cardiomyocyte hypertrophy, diastolic dysfunction, and cardiac remodeling relevant to HFpEF^5^. However, most studies using this model have focused on established or later-stage disease, after structural and functional abnormalities are already present. These end-stage analyses may overlook the initiating cellular events and communication networks that accompany disease onset and progression. Identifying these early events could reveal stage-specific mechanisms amenable to intervention before advanced pathological remodeling becomes established.

HFpEF progression involves multiple cardiac cell populations. Non-CM populations, including endothelial cells (ECs), fibroblasts (FBs), immune cells, smooth muscle cells (SMCs), and other stromal cells, are increasingly recognized as active regulators of the HFpEF microenvironment^6^. Recent studies support the concept that HFpEF involves coordinated remodeling of endothelial, immune, and stromal compartments, including coronary microvascular endothelial inflammation, immune activation, and FB-mediated extracellular matrix (ECM) remodeling^7–10^. However, the temporal and cell-type-specific programs that arise during HFpEF onset and progression, and how these compartments communicate over time, remain poorly defined.

Single-cell RNA sequencing provides the resolution needed to define disease-associated transcriptional programs at cell-type and subpopulation levels. However, existing single-cell studies of HFpEF have largely focused on single time points^7,11^, limiting their ability to capture dynamic cellular transitions that occur during disease progression. To address this gap, we applied time-series scRNA-seq to the HFD+L-NAME mouse HFpEF model, generating a longitudinal atlas of cardiac non-CMs across baseline (Ctrl/0w), early (1w), intermediate (4w), and established (8w) disease stages. By integrating scRNA-seq with metabolic, hemodynamic, echocardiographic, single-cardiomyocyte functional, and whole-heart bulk RNA-seq analyses, we aligned physiological disease progression with cell-type-specific transcriptional remodeling. We further assessed conservation of key endothelial programs in an L-NAME-independent HFpEF model and in human HFpEF single-nucleus RNA-seq (snRNA-seq) data and used an in vitro endothelial–macrophage adhesion assay to test whether HFpEF-like stress promotes endothelial inflammatory activation and macrophage adhesion. This integrated strategy defines the temporal vascular, immune, and stromal programs that accompany HFpEF progression and identifies candidate cell-type-specific and intercellular signaling mechanisms for future functional and therapeutic investigation.

## Results

### Longitudinal phenotyping reveals early and progressive systemic, cardiac, and molecular remodeling in HFD+L-NAME HFpEF

Most single-cell studies of the HFpEF model have focused on established or late-stage disease, often after prolonged metabolic and nitric oxide synthase–inhibitory stress, leaving the early and stage-dependent cellular programs that accompany disease onset and progression incompletely defined. To address this gap, we performed systematic longitudinal phenotyping and transcriptomic profiling using a two-hit mouse model induced by high-fat diet (HFD) combined with the nitric oxide synthase inhibitor Nω-nitro-L-arginine methyl ester hydrochloride (L-NAME), hereafter referred to as the HFD+L-NAME model. Mice maintained on normal chow served as controls. Metabolic phenotyping, blood pressure measurement, exercise testing, echocardiography, single-cardiomyocyte functional analysis, whole-heart bulk RNA-seq, and cardiac non-cardiomyocyte scRNA-seq were performed across baseline/control and sequential disease stages, corresponding to Ctrl/0w, early 1w, intermediate 4w, and established 8w disease after HFpEF induction (**Fig. 1A**).

**Figure 1.**
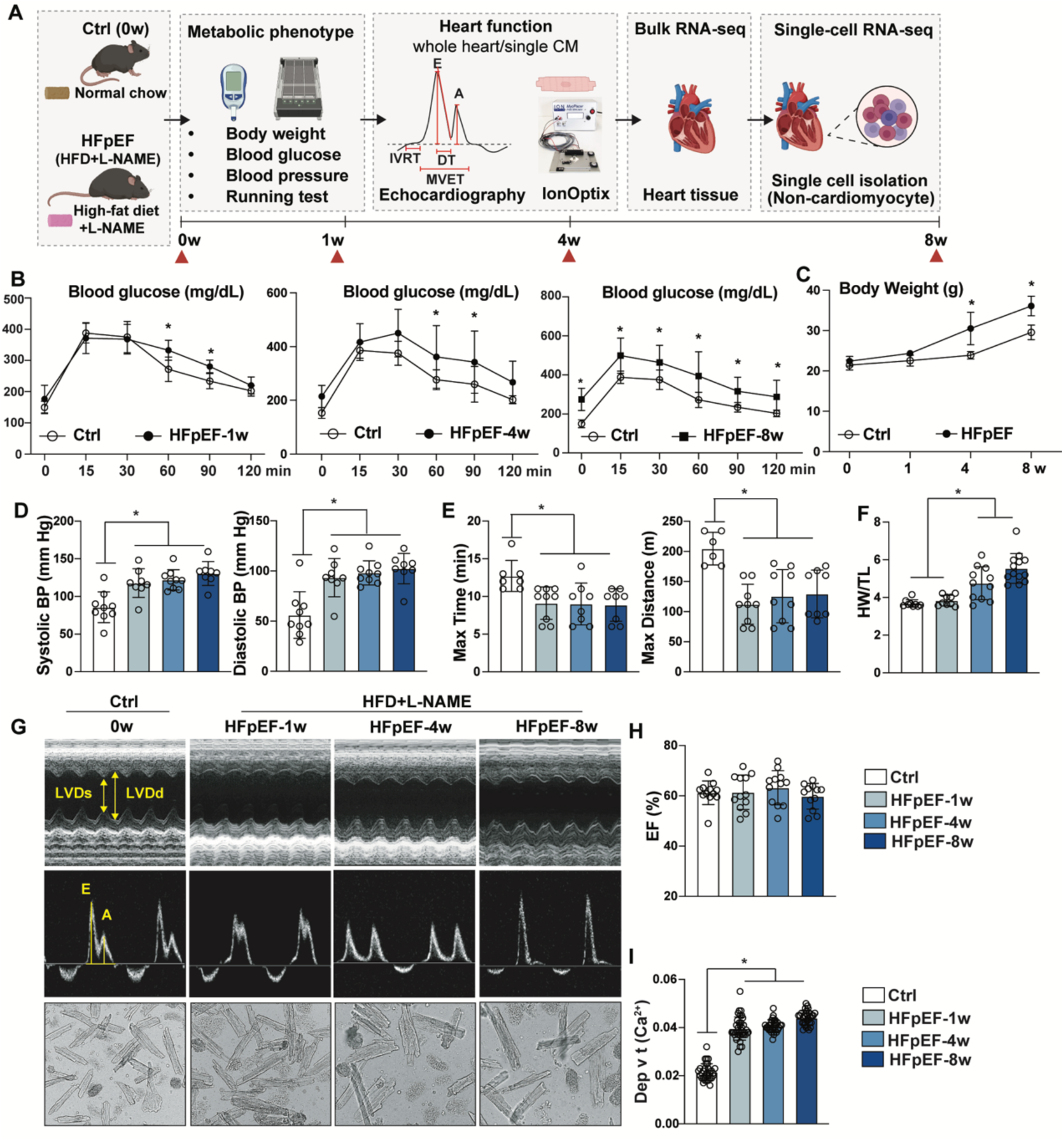
Longitudinal phenotype characterization and transcriptomic profiling of the metabolically stressed HFpEF models. **(A)** Schematic of longitudinal HFD+L-NAME HFpEF study design. Mice were maintained on normal control chow or HFD plus L-NAME and analyzed at baseline/0 week (Ctrl/0w), 1 week (1w), 4 weeks (4w), and 8 weeks (8w). Metabolic phenotyping, blood pressure measurement, running test, and echocardiography, single-cardiomyocyte functional assessment, bulk RNA-seq, and non-cardiomyocyte (non-CM) scRNA-seq were performed across the time course. **(B)** Glucose tolerance testing was measured across disease progression (Ctrl n=5, 1w n=9, 4w n=9, 8w n=7). Statistical analysis was performed using two-way ANOVA followed by Tukey’s multiple comparisons test. Data are presented as mean ± SEM. *P < 0.05 vs. Ctrl or indicated comparison. **(C)** Body weight was measured across disease progression (Ctrl n=10, 1w n=9, 4w n=10, 8w n=11). Statistical analysis was performed using two-way ANOVA followed by Tukey’s multiple comparisons test. Data are presented as mean ± SEM. *P < 0.05 vs. Ctrl or indicated comparison. **(D)** Systolic and diastolic blood pressure were measured across disease progression (Ctrl n=9, 1w n=8, 4w n=9, 8w n=8). Statistical analysis was performed using one-way ANOVA followed by Tukey’s multiple comparisons test. Data are presented as mean ± SEM. *P < 0.05 vs. Ctrl or indicated comparison. **(E)** Exercise capacity, assessed by maximal running time and maximal running distance, was measured across disease progression (Ctrl n=7, 1w n=8, 4w n=8, 8w n=8). Statistical analysis was performed using one-way ANOVA followed by Tukey’s multiple comparisons test. Data are presented as mean ± SEM. *P < 0.05 vs. Ctrl or indicated comparison. **(F)** Heart weight normalized to tibial length (HW/TL) was measured across disease progression Ctrl n=9, 1w n=9, 4w n=11, 8w n=13). Statistical analysis was performed using one-way ANOVA followed by Tukey’s multiple comparisons test. Data are presented as mean ± SEM. *P < 0.05 vs. control or indicated comparison. **(G)** Echocardiographic analysis of cardiac systolic and diastolic function across disease progression. **(H)** Quantification of echocardiographic parameters, including ejection fraction across disease progression (n=12 for each group). Statistical analysis was performed using one-way ANOVA followed by Tukey’s multiple comparisons test. Data are presented as mean ± SEM. *P < 0.05 vs. control or indicated comparison. **(I)** Single-cardiomyocyte calcium-handling measurement by IonOptix analysis. Statistical analysis was performed using one-way ANOVA followed by Tukey’s multiple comparisons test. (Ctrl n=31, 1w n=36, 4w n=33, 8w n=31). Data are presented as mean ± SEM. *P < 0.05 vs. control or indicated comparison.

Longitudinal phenotyping showed that systemic abnormalities were detectable at the earliest profiled disease stage. Glucose tolerance testing revealed impaired glucose clearance in HFD+L-NAME mice, with elevated post-challenge blood glucose detectable by 1w and persisting through 4w and 8w (**Fig. 1B**). Body weight increased progressively at 4w and 8w but was not markedly elevated at 1w (**Fig. 1C**), indicating that metabolic dysfunction was detectable before overt weight gain became prominent. Systolic and diastolic blood pressure were elevated in HFD+L-NAME mice compared with controls across the disease course (**Fig. 1D**). Exercise testing further demonstrated reduced maximal running time and maximal running distance at the profiled stages, indicating reduced exercise tolerance beginning early in disease progression (**Fig. 1E**). Together, these data show that impaired glucose handling, elevated blood pressure, and reduced exercise tolerance are already present at early disease stages and persist through established disease. Cardiac structural and functional abnormalities were assessed across the same longitudinal time course. Heart weight normalized to tibial length increased most prominently at 4w and 8w, consistent with progressive cardiac hypertrophic remodeling (**Fig. 1F**). Echocardiographic analysis showed preserved ejection fraction throughout the disease course, consistent with maintained systolic function at the profiled stages (**Fig. 1G and H).** In contrast, Doppler echocardiography revealed altered transmitral filling profiles beginning at early disease stages and persisting over time, consistent with abnormal diastolic filling in the setting of preserved ejection fraction (**Fig. 1G**). At the single-CM level, isolated CMs from HFD+L-NAME hearts demonstrated altered calcium-handling kinetics by IonOptix analysis **(Fig. 1G and I)**, suggesting that cell-intrinsic CM functional abnormalities accompany the early disease course.

To determine whether these phenotypic abnormalities were accompanied by coordinated molecular remodeling, we performed whole-heart bulk RNA-seq across an extended time course that included Ctrl/0w, 1w, 4w, 8w, and 12w. Principal component analysis (PCA) separated control and HFD+L-NAME hearts and further distinguished disease stages along the disease-time axis, indicating progressive and time-dependent transcriptomic remodeling (**Supplementary Fig. 1A**). Differentially expressed gene (DEG) analysis identified substantial transcriptional changes between HFD+L-NAME and control hearts, as well as stage-dependent transitions during disease progression (**Supplementary Fig. 1B**). Notably, the 8w versus 12w comparison revealed only modest additional transcriptomic differences, with 42 upregulated and 7 downregulated genes, indicating that major whole-heart molecular remodeling was largely established by 8w with limited additional transcriptomic change thereafter. Pathway analysis of genes associated with the dominant principal components revealed that the major sources of transcriptomic variation included translation, peptide metabolism, mitochondrial respiratory chain complex assembly, angiogenesis, immune system processes, RNA processing, mitochondrial translation, actin-based movement, cardiac muscle cell action potential, and cell migration/motility pathways (**Supplementary Fig. 1C**). These pathway-level findings indicate that whole-heart molecular remodeling involves metabolic, mitochondrial, immune, vascular, and structural programs that emerge early and evolve across the disease course.

Together, this longitudinal phenotyping and bulk RNA-seq data demonstrate that systemic, cardiac, and molecular abnormalities are detectable by 1w and persist or progress through 4w and 8w. Because whole-heart bulk RNA-seq cannot identify the specific cellular sources of tissue-level remodeling, single-cell resolution was necessary to define the cell types driving these molecular changes^12^. We next performed scRNA-seq of cardiac non-CMs to define the vascular, stromal, and immune programs underlying HFpEF progression at cellular resolution. Because the 8w stage captured established disease with limited additional whole-heart transcriptomic change by 12w, we selected Ctrl/0w, 1w, 4w, and 8w for downstream scRNA-seq. This design enabled resolution of baseline, early, intermediate, and established cellular remodeling programs across disease progression rather than focusing on a single late-stage endpoint.

### Cellular landscape of cardiac non-CMs during HFpEF progression revealed by scRNA-seq

To define cell-type-specific remodeling of the cardiac non-CM compartment during HFpEF progression, we performed scRNA-seq on non-CMs isolated from Ctrl/0w and HFD+L-NAME HFpEF hearts at 1w, 4w, and 8w (**Fig. 2A**). After quality control and filtering, 94,848 high-quality cardiac non-CMs were retained for downstream analysis (**Supplementary Fig. 2A-C**). Unsupervised clustering and UMAP visualization resolved nine major cardiac non-CM populations across the time course, including endothelial cells (ECs), fibroblasts (FBs), smooth muscle cells (SMCs), macrophages and mononuclear phagocytes (MPs), T cells (T), B cells (B), neutrophils (NEUTs), natural killer cells (NKs), and platelets (PLTs) (**Fig. 2B**). Canonical marker analysis confirmed the identity of each population, with enrichment of *Pecam1*, *Kdr*, *Cdh5*, and *Vwf* in ECs; *Col1a1*, *Col3a1*, and *Dcn* in FBs; *Acta2*, *Tagln*, and *Myh11* in SMCs; *C1qa*, *Lyz2*, *Csf1r*, and *Apoe* in MPs; *Cd3d*, *Trac*, and *Il7r* in T cells; *Cd79a*, *Ms4a1*, and *Cd19* in B cells; *S100a8*, *S100a9*, and *Retnlg* in NEUTs; *Nkg7*, *Ncr1*, *Klrk1*, *Gzmb*, and *Prf1* in NK cells; and *Pf4*, *Ppbp*, and *Itga2b* in PLTs (**Fig. 2C**). These lineage-specific marker patterns confirmed successful integration and annotation of the major vascular, stromal, and immune populations in the cardiac non-CM compartment.

**Figure 2.**
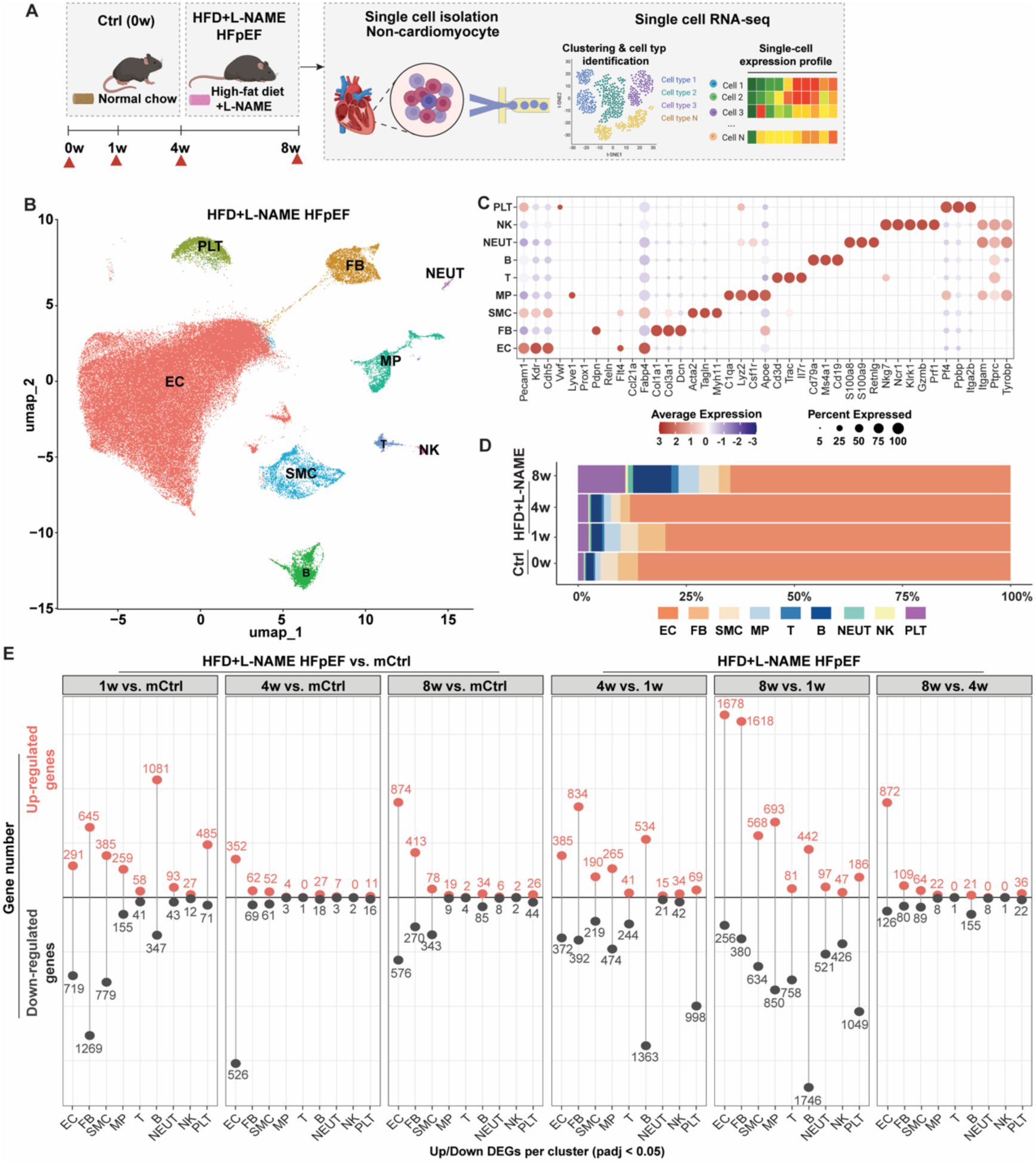
Single-cell profiling of non-CMs during HFpEF progression. **(A)** Schematic of non-CM isolation and scRNA-seq workflow from control and HFD+L-NAME HFpEF hearts at 1, 4, and 8 weeks. **(B)** Uniform manifold approximation and projection (UMAP) visualization of 94,848 cardiac non-CMs, identifying endothelial cells (EC), fibroblasts (FB), smooth muscle cells (SMC), macrophages (MP), T cells, B cells, neutrophils (NEUT), NK cells, and platelets (PLT). **(C)** Dot plot showing canonical marker gene expression used for cell-type annotation. Dot size represents the percentage of cells expressing each marker, and color represents scaled average expression. **(D)** Proportion plot showing relative abundance of non-CM populations. **(E)** Lollipop plot showing upregulated and downregulated differentially expressed genes (DEGs) across major cell populations and disease-stage comparisons. DEGs were defined using adjusted P < 0.05 and log fold change > 0.25.

To further support cell-type annotation at the pathway level, we performed GO enrichment analysis using cell-type-enriched genes (**Supplementary Fig. 2D**). EC-enriched genes were associated with regulation of vasculature development, regulation of angiogenesis, EC migration, and EC proliferation. FB-enriched genes were associated with ECM organization, extracellular structure organization, and connective tissue development, whereas SMC-enriched genes were enriched for muscle system processes, muscle contraction, muscle cell differentiation, and calcium ion transport. Immune-cell populations showed lineage-consistent enrichment patterns: MPs and NEUTs were enriched for myeloid leukocyte activation, regulation of inflammatory response, chemotaxis, granulocyte migration, and responses to bacterial-origin molecules; T cells were enriched for lymphocyte differentiation, T-cell differentiation, T-cell proliferation, and T-cell receptor signaling; B cells were enriched for B-cell differentiation and immune-cell activation programs; and NK cells were enriched for cell killing and natural killer cell-mediated cytotoxicity. PLT-enriched genes were associated with platelet activation, blood coagulation, wound healing, and regulation of body fluid levels. These pathway-level signatures were concordant with canonical cell-type biology, providing independent support for the major cell-type assignments and indicating that the scRNA-seq dataset captured the major vascular, stromal, immune, and platelet compartments of the cardiac non-cardiomyocyte landscape during HFpEF progression.

Cell composition analysis showed that ECs represented the most abundantly recovered non-CM population, followed by FBs and SMCs (**Fig. 2D**). Although all major populations were detected across control and HFD+L-NAME hearts, their relative abundance changed over time. Notably, the relative proportion of ECs decreased at 8w, whereas B cells and PLTs increased at this later stage (**Fig. 2D**). These changes suggest progressive remodeling of the cardiac stromal, vascular, and immune microenvironment during HFpEF progression, while recognizing that scRNA-seq composition reflects both biological abundance and cell-recovery efficiency. To assess transcriptional remodeling within each population, we next identified differentially expressed genes (DEGs) across pairwise disease-stage comparisons using an adjusted P value < 0.05 and log fold change > 0.25 (**Fig. 2E**). Substantial transcriptional changes were already present at 1w across multiple cell types, indicating that the cardiac non-CM compartment responds early to HFD+L-NAME stress. Across the time course, ECs and FBs showed particularly extensive transcriptional remodeling, especially in comparisons involving later disease stages and stage-to-stage transitions. ECs displayed robust transcriptional remodeling in HFpEF versus control comparisons and remained highly dynamic during disease progression, supporting ECs as a major disease-responsive population. FBs also exhibited extensive transcriptional changes, consistent with progressive stromal remodeling during established disease. These findings indicate that HFpEF progression is accompanied by broad but cell-type-selective remodeling, with vascular and stromal compartments representing dominant components of the evolving non-CM response.

Together, these data establish a time-resolved single-cell atlas of the cardiac non-CM compartment during HFD+L-NAME HFpEF progression. The predominance of ECs among recovered cells, together with their strong and dynamic transcriptional response across disease stages, identified the endothelial compartment as a major candidate driver of early cardiac microenvironmental remodeling. We therefore next focused on endothelial subclustering to define disease-associated endothelial activation states and their potential contribution to HFpEF progression.

### Endothelial reclustering reveals heterogeneous and dynamically activated EC states during HFpEF progression

The global non-CM atlas identified that ECs were the predominant recovered non-CM population and a highly disease-responsive compartment during HFpEF progression (**Fig. 2**). Because endothelial dysfunction and microvascular inflammation are central features of HFpEF pathobiology, we next focused on the endothelial population to define disease-associated EC heterogeneity and activation at subpopulation resolution. We reclustered ECs from the integrated scRNA-seq dataset and identified transcriptionally distinct EC subpopulations, designated EC-1 through EC-8 (**Fig. 3A**). All EC subclusters expressed canonical endothelial markers, including *Pecam1*, *Cdh5*, and *Kdr*, confirming their endothelial identity (**Fig. 2C and Fig. 3B**). Subcluster annotation using established marker genes and functional transcriptional signatures revealed distinct EC states, including capillary ECs, stress-response ECs, arterial ECs, interferon-stimulated ECs, inflammatory ECs, migratory ECs, ECM-remodeling ECs, and proliferating ECs (**Fig. 3B**). Capillary EC subsets were characterized by *Aqp1*, *Aplnr*, *Car4*, and *Car8*; stress-response genes such as *Junb*, *Fos*, *Hspa1a*, and *Atf3*; arterial ECs marked by *Hey1*, *Sema3g*, *Unc5b*, and *Gja5*; interferon-stimulated ECs marked by *Ifit3*, *Isg15*, *Irf7*, *Stat1*, and *Stat2*^13^; inflammatory ECs marked by *Nr2f2*, *Ptgs1*, *Vcam1*, and *Il1r1*; migratory ECs marked by *Marcksl1*, *Rhoc*, and *Dpysl3*; ECM-remodeling ECs marked by *Fbln5*, *Col18a1*, and *Fn1*; and proliferating ECs marked by *Mki67*, *Top2a*, *Cdk1*, and *Ccna2*^14^. Together, these findings indicate that cardiac ECs are transcriptionally heterogeneous and comprise multiple specialized vascular, inflammatory, remodeling, and proliferative states during HFpEF progression.

**Figure 3.**
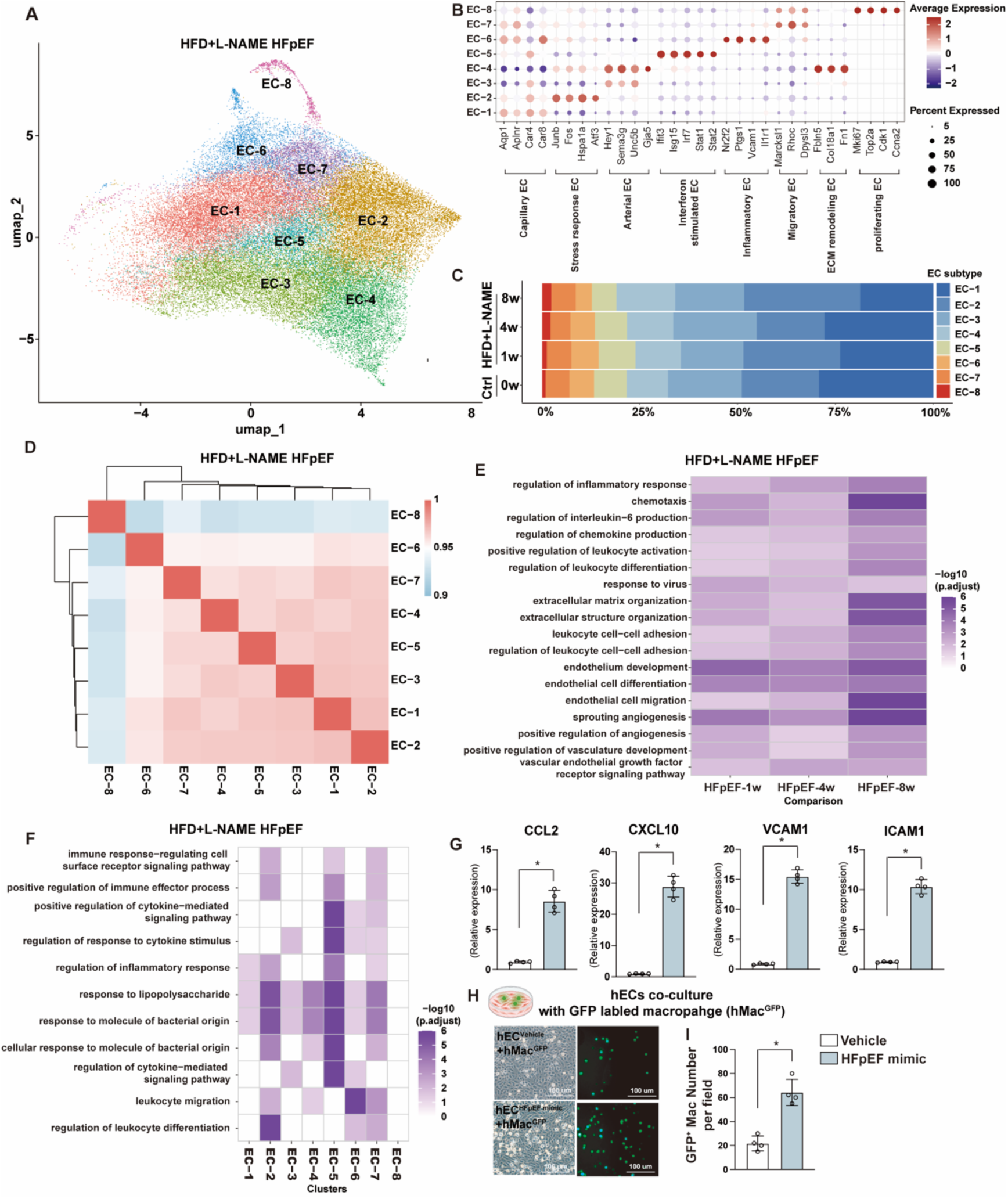
Endothelial cell heterogeneity and disease-associated activation programs in HFD+L-NAME HFpEF. **(A)** UMAP visualization of reclustered endothelial cells, identifying eight endothelial subtypes (EC-1 through EC-8). **(B)** Dot plot showing signature genes used to annotate endothelial subtypes, including capillary EC, stress-response EC, arterial EC, interferon-stimulated EC, inflammatory EC, migratory EC, ECM-remodeling EC, and proliferating EC. Dot size indicates the percentage of cells expressing each marker, and color indicates scaled average expression. **(C)** Proportion plot showing relative abundance of endothelial subtypes across control and HFpEF stages. **(D)** Heatmap showing pairwise transcriptional similarity of eight EC subpopulations by Spearman correlation analysis. **(E)** GO enrichment analysis of disease-upregulated genes in ECs across HFpEF progression, highlighting inflammatory response, chemotaxis, cytokine signaling, leukocyte adhesion, endothelial migration, angiogenesis, ECM organization, and VEGF receptor signaling pathways. Color intensity represents −log10(adjusted P-val). **(F)** Cluster-level GO enrichment analysis across EC subpopulations, showing subtype-specific immune, inflammatory, cytokine-response, and leukocyte-interaction programs. Color intensity represents −log10(adjusted P-val). **(G)** Expression of endothelial activation and adhesion genes, including CXCL10, CCL2, ICAM1, and VCAM1, in human endothelial cells (hECs) exposed to HFpEF-like metabolic and inflammatory stress (HFpEF mimic) versus vehicle-treated controls (vehicle). Statistical analysis was performed using a paired t-test. (n=4). Data are presented as mean ± SEM. *P < 0.05 vs. vehicle. **(H)** Representative phase-contrast and fluorescence microscopy images of GFP-labeled human macrophage (hMac^GFP^) adhesion to confluent human endothelial (hEC) monolayers. **(I)** Quantification of macrophage (hMac^GFP^) adhesion to endothelial monolayers (hECs) under HFpEF-like stress (HFpEF mimic) versus vehicle-treated control conditions (Vehicle). Statistical analysis was performed using a paired t-test. (n=4). Data are presented as mean ± SEM. *P < 0.05 vs. vehicle.

The relative abundance of EC subtypes changed across disease stages, indicating dynamic remodeling of the endothelial compartment (**Fig. 3C**). Pairwise transcriptional similarity analysis showed that most EC subpopulations shared related endothelial transcriptional programs, whereas EC-8 was more transcriptionally distinct, consistent with its proliferative identity and enrichment of cell-cycle genes (**Fig. 3D**). These data suggest that HFpEF progression reshapes the endothelial landscape through both shifts in EC subtype composition and emergence of specialized activation states.

To define disease-associated endothelial programs, we performed differential expression and GO enrichment analyses of genes upregulated in HFpEF ECs at each disease stage compared with controls (**Fig. 3E**). HFpEF ECs were enriched for several categories of biological process. Inflammatory and chemokine-related programs included regulation of inflammatory response, regulation of interleukin-6 production, regulation of chemokine production, and chemotaxis. Leukocyte-interaction programs included positive regulation of leukocyte activation, leukocyte cell-cell adhesion, and regulation of leukocyte differentiation, suggesting increased endothelial capacity to engage immune cells. Vascular remodeling programs included endothelial cell migration, sprouting angiogenesis, positive regulation of angiogenesis, endothelium development, and VEGF receptor signaling. These enrichment patterns were already detectable at 1w and became more pronounced at later stages, indicating that inflammatory, immune-interaction, and vascular remodeling programs are early and sustained components of the HFpEF endothelial response.

Cluster-level pathway analysis further showed that these inflammatory and immune-interaction programs were not uniformly distributed across all ECs but were concentrated within specific endothelial states (**Fig. 3F**). Multiple endothelial subclusters were enriched for immune response-regulating cell-surface receptor signaling, cytokine-mediated signaling, regulation of inflammatory response, response to lipopolysaccharide and bacterial-origin molecules, leukocyte migration, and regulation of leukocyte differentiation. Among these, EC-5 showed the most prominent enrichment of cytokine-mediated signaling, inflammatory response, and immune effector-related pathways, identifying it as a highly activated inflammatory endothelial state. EC-6 showed strong enrichment for leukocyte migration-related programs, whereas EC-2 was enriched for immune effector and leukocyte differentiation pathways. EC-7 shared features of both programs, with enrichment for leukocyte migration as well as immune effector/leukocyte differentiation pathways. These subtype-level differences indicate that endothelial inflammatory activation in HFpEF is organized across specialized endothelial states rather than occurring as a uniform pan-endothelial response.

To functionally evaluate the scRNA-seq prediction that HFpEF-associated stress promotes an activated endothelial state with enhanced immune-interacting capacity, we exposed human endothelial cells (hECs) to combined metabolic and inflammatory stress designed to mimic HFpEF-like conditions, with vehicle-treated cells serving as controls. HFpEF-mimic stress significantly increased expression of endothelial activation, adhesion, and chemokine genes, including CCL2, CXCL10, VCAM1, and ICAM1 (**Fig. 3G**). To determine whether this transcriptional activation was associated with increased macrophage adhesion, vehicle- or HFpEF-mimic-stressed hEC monolayers were co-cultured with GFP-labeled human macrophages. HFpEF-mimic-stressed endothelial monolayers demonstrated significantly greater macrophage adhesion than vehicle-treated controls, as quantified by GFP-positive macrophage number per field (**Fig. 3H**).

Together, these results demonstrate that HFpEF progression is accompanied by dynamic endothelial heterogeneity and coordinated activation of specialized EC subpopulations. The endothelial response encompasses inflammatory signaling, interferon-associated activation, leukocyte-interaction programs, vascular remodeling, and proliferative adaptation. The convergence of time-resolved scRNA-seq with functional endothelial validation supports endothelial activation as an early and prominent feature of cardiac non-CM remodeling during HFpEF progression and provides a foundation for subsequent analysis of endothelial–immune communication networks.

### Cross-model analysis supports conserved endothelial activation across mechanistically distinct HFpEF models

The HFD+L-NAME model combines metabolic stress with nitric oxide synthase inhibition; therefore, L-NAME may directly affect nitric oxide signaling and influence endothelial phenotypes. To determine whether the endothelial activation programs identified in the HFD+L-NAME model were conserved in a L-NAME-independent HFpEF context, we analyzed a second, mechanistically distinct HFpEF model induced by HFD combined with mild transverse aortic constriction (mTAC), referred to here as HFD+mTAC^15^. This model combines metabolic stress with mild mechanical pressure overload and has been shown to induce diastolic dysfunction with preserved systolic function, metabolic abnormalities, and cardiac hypertrophic remodeling in the absence of L-NAME exposure. We used the HFD+mTAC dataset as a focused cross-model validation cohort for endothelial remodeling rather than as a second complete cell-type-resolved atlas.

Cardiac non-CMs were isolated from sham/0w and HFD+mTAC hearts at 1w, 4w, and 8w stages and profiled by scRNA-seq (**Fig. 4A**). Cells were filtered to retain unique feature counts between 200 and 3,000, and doublets were removed using a consensus approach combining a density-based algorithm (computeDoubletDensity) and a KNN-based classifier (scDblFinder) (**Supplementary Fig. 3A**). Quality-control metrics (nFeature_RNA, nCount_RNA, percent.mt) were consistent across all four timepoints (**Supplementary Fig. 3B**). Batch effects across timepoints were assessed and corrected using Harmony prior to unbiased clustering and cell-type annotation. UMAP embedding following batch correction showed extensive intermixing of cells from sham, 1w, 4w, and 8w timepoints across all major clusters, indicating effective integration without strong timepoint-driven batch artifacts (**Supplementary Fig. 3C**). Unsupervised clustering identified the major cardiac non-CM populations, including ECs, FBs, SMCs, MPs, T, B, NEUTs, and PLTs (**Supplementary Fig. 3D**). Gene Ontology enrichment analysis of cluster-defining marker genes confirmed expected biological identities for each population, including vasculature development and angiogenesis-related terms in EC, ECM and connective tissue organization in FB, muscle contraction and calcium ion transport in SMC, leukocyte activation and chemotaxis programs in MP, T cells, and B cells, neutrophil chemotaxis and migration in NEUT, and blood coagulation and wound healing in PLT (**Supplementary Fig. 3E**), supporting the accuracy of cell-type annotation in the HFD+mTAC dataset. Similar to the HFD+L-NAME model, ECs represented a major recovered non-CM population across the HFD+mTAC time course, with additional representation of stromal and immune populations (**Fig. 4B and C**). Differential expression analysis further revealed dynamic, cell-type-specific transcriptional remodeling across disease stages, confirming that the HFD+mTAC model induces broad remodeling of the cardiac non-CM compartment (**Supplementary Fig. 4A**).

**Figure 4.**
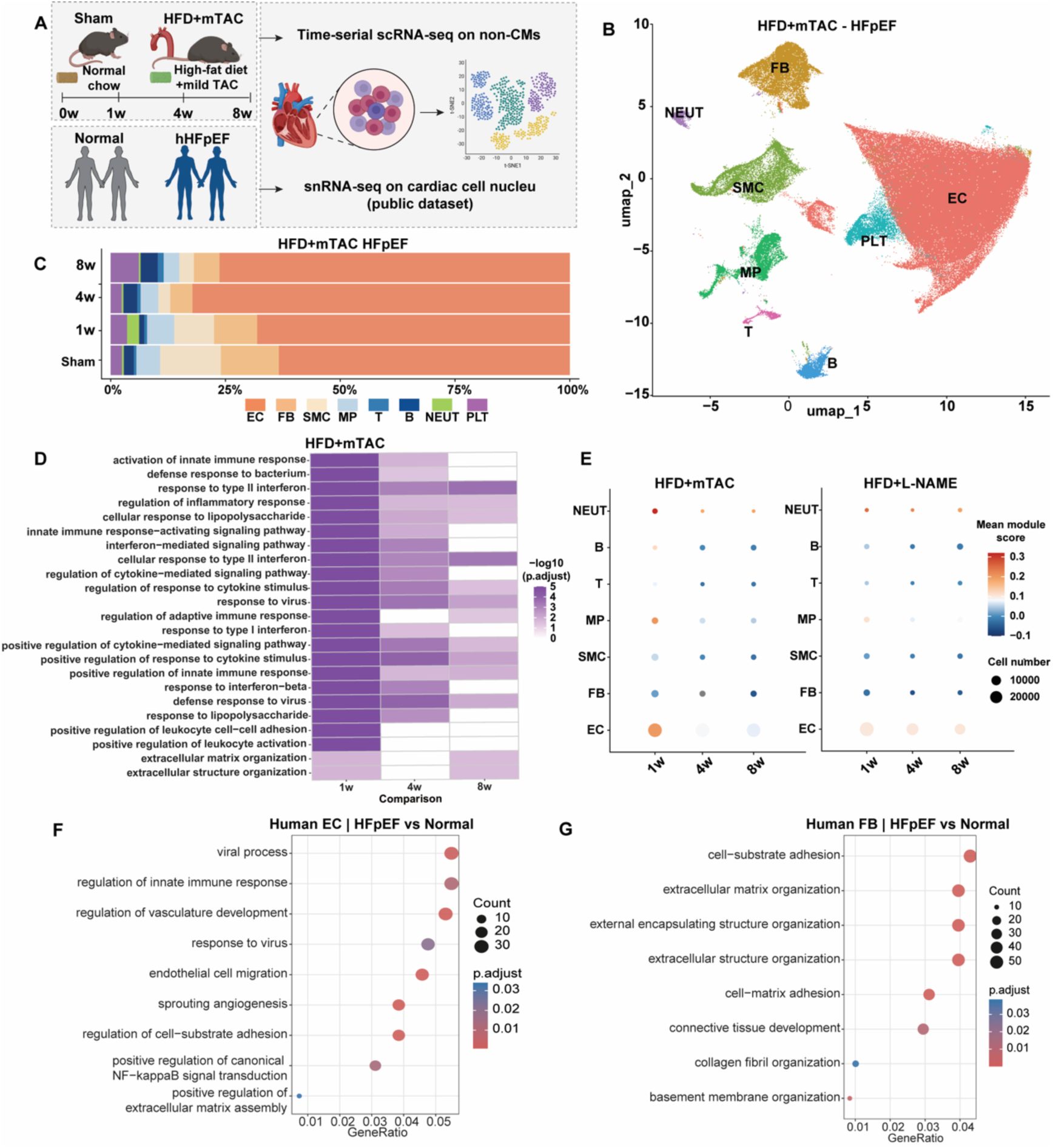
Endothelial activation is conserved across mechanistically distinct HFpEF models and human disease. **(A)** Schematic comparison of the HFD+L-NAME and HFD+mTAC HFpEF models. The HFD+L-NAME model combines metabolic stress with nitric oxide synthase inhibition, whereas the HFD+mTAC model combines metabolic stress with mild mechanical pressure overload without L-NAME exposure. Cardiac non-CMs were isolated from HFD+mTAC hearts at sham (0 week), 1 week (1w), 4 weeks (4w), and 8 weeks (8w) for scRNA-seq. **(B)** UMAP visualization of non-CMs from HFD+mTAC hearts, identifying EC, FB, SMC, MP, T cells, B cells, NEUT, and PLT populations. **(C)** Proportion plot showing relative abundance of non-CM populations across sham and HFpEF-like stages. **(D)** GO enrichment analysis of disease-upregulated genes in ECs from HFD+mTAC hearts across disease progression. Color intensity represents −log10(adjusted P-val). HFD+mTAC ECs showed prominent early enrichment of innate immune, interferon-response, inflammatory-response, leukocyte-interaction, and extracellular structure organization pathways. **(E)** Module scoring of an EC-derived disease-response module across major non-CM populations. This module excluded canonical endothelial identity markers and comprised inducible inflammatory, adhesion, chemokine, and interferon-response genes. Dot size represents cell number, and color represents mean module score. **(F)** GO enrichment analysis of disease-associated genes in human HFpEF endothelial cells, highlighting innate immune response, viral and interferon-associated response, regulation of vasculature development, endothelial migration, sprouting angiogenesis, regulation of cell-substrate adhesion, NF-κB-related signaling, and ECM assembly. **(G)** GO enrichment analysis of disease-associated genes in human HFpEF fibroblasts, highlighting cell-substrate adhesion, ECM organization, external encapsulating structure organization, extracellular structure organization, cell-matrix adhesion, connective tissue development, collagen fibril organization, and basement membrane organization.

We then focused on endothelial genes upregulated in HFD+mTAC hearts relative to sham controls. GO enrichment analysis showed that HFD+mTAC ECs were prominently enriched for innate immune and inflammatory pathways at 1w, including activation of innate immune response, response to type II interferon, regulation of inflammatory response, cellular response to lipopolysaccharide, interferon-mediated signaling, response to viral and bacterial-origin molecules, leukocyte activation, and ECM organization (**Fig. 4D**). These programs were most prominent at the early disease stage and became less uniformly enriched at 4w and 8w, suggesting that endothelial inflammatory activation in the HFD+mTAC model is early and dynamic rather than simply progressive. The presence of overlapping inflammatory, interferon-response, leukocyte-interaction, and ECM-remodeling pathways in this L-NAME-independent model supports the idea that endothelial activation is not solely a consequence of nitric oxide synthase inhibition.

To compare endothelial disease programs across models, we scored an endothelial disease-response module composed of inducible inflammatory, adhesion, chemokine, and interferon-response genes, while excluding canonical endothelial identity markers (**Fig. 4E).** In both HFD+mTAC and HFD+L-NAME hearts, the module was preferentially enriched in ECs compared with most other non-CM populations, with the strongest increase occurring at early disease stages. These findings indicate that the endothelial activation signature identified in the HFD+L-NAME model is reproduced in an independent HFpEF model and reflects a conserved endothelial-enriched disease response rather than a model-specific effect of L-NAME exposure.

To assess the translational relevance of these mouse-derived endothelial programs, we further analyzed public human HFpEF single-nucleus RNA-seq data (snRNA-seq)^10^. The human dataset resolved major cardiac cell populations, including EC (EC1, EC2, and Endocardial EC (EndoC), FB, pericytes, vascular smooth muscle cells (VSMC), cardiomyocytes (CM), MP, lymphoid cells (LYM), mast cells (Mast), neuronal cells (Neur), and adipocytes (Adipo), with cell identities supported by canonical marker expression (**Supplementary Fig. 4B–D**). GO enrichment analysis of human HFpEF ECs identified pathways related to innate immune and antiviral responses, regulation of vascular development, EC migration, sprouting angiogenesis, regulation of cell-substrate adhesion, NF-κB signaling, and ECM assembly (**Fig. 4F**). These pathways paralleled the inflammatory, adhesion-associated, and vascular-remodeling programs observed in both mouse HFpEF models. In parallel, human HFpEF FBs were enriched for ECM organization, cell-substrate adhesion, cell-matrix adhesion, connective tissue development, collagen fibril organization, and basement membrane organization (**Fig. 4G**), consistent with stromal remodeling as a conserved component of the HFpEF non-CM response.

Together, these cross-model and human single-nucleus analyses support conservation of endothelial inflammatory, interferon-response, adhesion-associated, and vascular-remodeling programs across mechanistically distinct HFpEF contexts. The replication of this endothelial activation signature in an L-NAME-independent mouse model and its alignment with human HFpEF endothelial programs strengthen the conclusion that endothelial activation represents a conserved feature of HFpEF-associated cardiac microvascular remodeling.

### Fibroblast subclustering reveals stage-dependent matrix remodeling and immune-stromal programs during HFpEF progression

Having established that ECs undergo inflammatory and immune-interacting activation during HFpEF progression, we next examined FBs, another major non-CM population implicated in ECM remodeling, tissue stiffness, and stromal signaling. To define FB remodeling at higher resolution, we reclustered FBs from the integrated scRNA-seq dataset. FBs were identified by robust expression of canonical FB markers, including Dcn, Col1a1, and Col1a2^7^ (**Fig. 5B**). Reclustering analysis revealed three distinct FB subpopulations, designated FB-1 through FB-3 **(Fig. 5A)**. FB-1 and FB-2 represented the major FB populations across control and HFpEF conditions, whereas FB-3 was a smaller subset (**Fig. 5C**).

**Figure 5.**
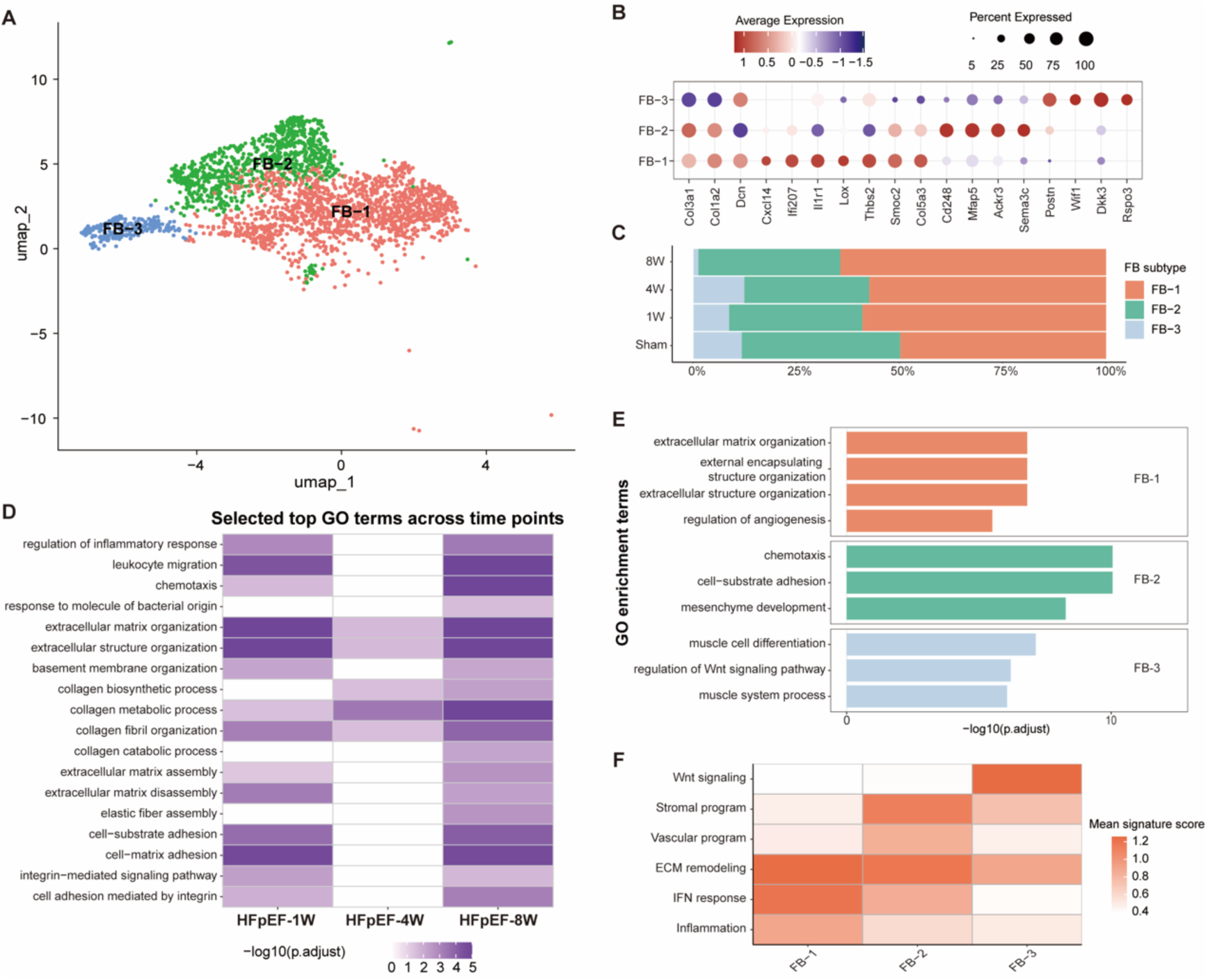
Fibroblast heterogeneity and stromal remodeling in HFD+L-NAME HFpEF. **(A)** UMAP visualization of reclustered fibroblasts, identifying three fibroblast subtypes (FB-1 through FB-3). **(B)** Dot plot showing marker genes associated with fibroblast subtypes. Dot size represents the percentage of cells expressing each marker, and color represents scaled average expression. **(C)** Proportion plot showing relative abundance of fibroblast subtypes across control and HFpEF stages. **(D)** GO enrichment analysis of disease-upregulated genes in fibroblasts from HFD+L-NAME hearts across HFpEF progression. Color intensity represents −log10(adjusted P-val). HFpEF fibroblasts showed enrichment of ECM organization, extracellular structure organization, basement membrane organization, collagen biosynthetic and metabolic processes, collagen fibril organization, ECM assembly and disassembly, elastic fiber assembly, cell-substrate adhesion, cell-matrix adhesion, and integrin-mediated signaling pathways. **(E)** Subtype-specific GO enrichment analysis showing functional specialization of FB-1, FB-2, and FB-3. Bar length represents −log10(adjusted P-value). **(F)** Heatmap of fibroblast gene program scores, including Wnt signaling, stromal program, vascular program, ECM remodeling, IFN response, and inflammation.

To characterize disease-associated FB activation, we performed differential expression and GO enrichment analyses across HFpEF stages. In contrast to the inflammatory and immune-interacting programs that dominated in ECs, FB responses were dominated by ECM and adhesion-related pathways. Genes upregulated in HFpEF FBs were enriched for ECM organization, extracellular structure organization, basement membrane organization, collagen biosynthetic and metabolic processes, collagen fibril organization, ECM assembly and disassembly, elastic fiber assembly, cell-substrate adhesion, cell-matrix adhesion, and integrin-mediated signaling pathways (**Fig. 5D**). These ECM-associated programs were most prominent at the late 8w stage, indicating that stromal matrix remodeling becomes increasingly pronounced during established HFpEF progression.

Inflammatory and immune-interacting pathways showed a stage-dependent pattern. Terms related to regulation of inflammatory response, leukocyte migration, chemotaxis, response to bacterial-origin molecules, cell-substrate adhesion, cell-matrix adhesion, and integrin-mediated signaling were enriched at the early 1w stage, less prominent at 4w, and re-enriched at 8w (**Fig. 5D**). This temporal pattern suggests that fibroblast remodeling during HFpEF progression may involve an early stress-associated inflammatory/adhesive response, followed by a later phase in which matrix remodeling and immune-stromal interaction programs occur together.

We next examined whether three FB subpopulations displayed distinct functional programs. Subtype-level pathway enrichment and gene-program scoring revealed functional specialization among FB-1, FB-2, and FB-3 (**Fig. 5E and F**). FB-1 was enriched for ECM organization, external encapsulating structure organization, extracellular structure organization, and regulation of angiogenesis, consistent with a matrix-remodeling and vascular-supportive FB state. FB-2 was enriched for chemotaxis, cell-substrate adhesion, and mesenchyme development, together with stromal and vascular-related programs, suggesting a role in microenvironment remodeling and immune-stromal communication. FB-3 showed enrichment of muscle cell differentiation, muscle system process, and regulation of Wnt signaling, consistent with a distinct remodeling-associated FB state (**Fig. 5E and F**).

Together, these findings demonstrate that FBs undergo dynamic and functionally heterogeneous remodeling during HFpEF progression. Whereas endothelial remodeling is characterized by inflammatory activation and immune-interaction programs, fibroblast remodeling is dominated by extracellular matrix organization, collagen remodeling, adhesion, and stromal signaling. These results suggest that FBs are key contributors to progressive matrix remodeling and stage-dependent immune-stromal remodeling within the HFpEF cardiac microenvironment.

### Macrophage reclustering reveals loss of resident-like macrophages and expansion of inflammatory and proliferative states during HFpEF progression

Because endothelial and FB programs both implicated inflammatory signaling, leukocyte interaction, and immune-stromal communication, we next examined macrophage/mononuclear phagocytes as a potential cellular bridge linking endothelial activation with FB-mediated stromal remodeling.

Macrophages/mononuclear phagocytes (MPs) were subsetted from the scRNA-seq dataset and reclustered, identifying four distinct MP subpopulations (MP-1 through MP-4) (**Fig. 6A**). MP-1 was enriched for resident-like macrophage markers, including *Lyve1*, *Folr2*, *Timd4*, *Mertk*, *Fcrls*, and *Cd163*, consistent with a homeostatic resident-like macrophage population (**Fig. 6B and C**). MP-2 expressed TREM-associated genes, including *Trem1*, *Trem3*, and *Treml4*, consistent with an inflammatory macrophage program. MP-3 was characterized by elevated *Ccr2* and *Trem1* expression, suggesting a highly inflammatory CCR2+TREM1+ macrophage state linked to immune-cell recruitment, inflammatory amplification, and adverse cardiac remodeling. MP-4 was distinguished by high expression of proliferation-associated genes, including *Mki67*, *Top2a*, and *Cdk1*, identifying a proliferating macrophage population.

**Figure 6.**
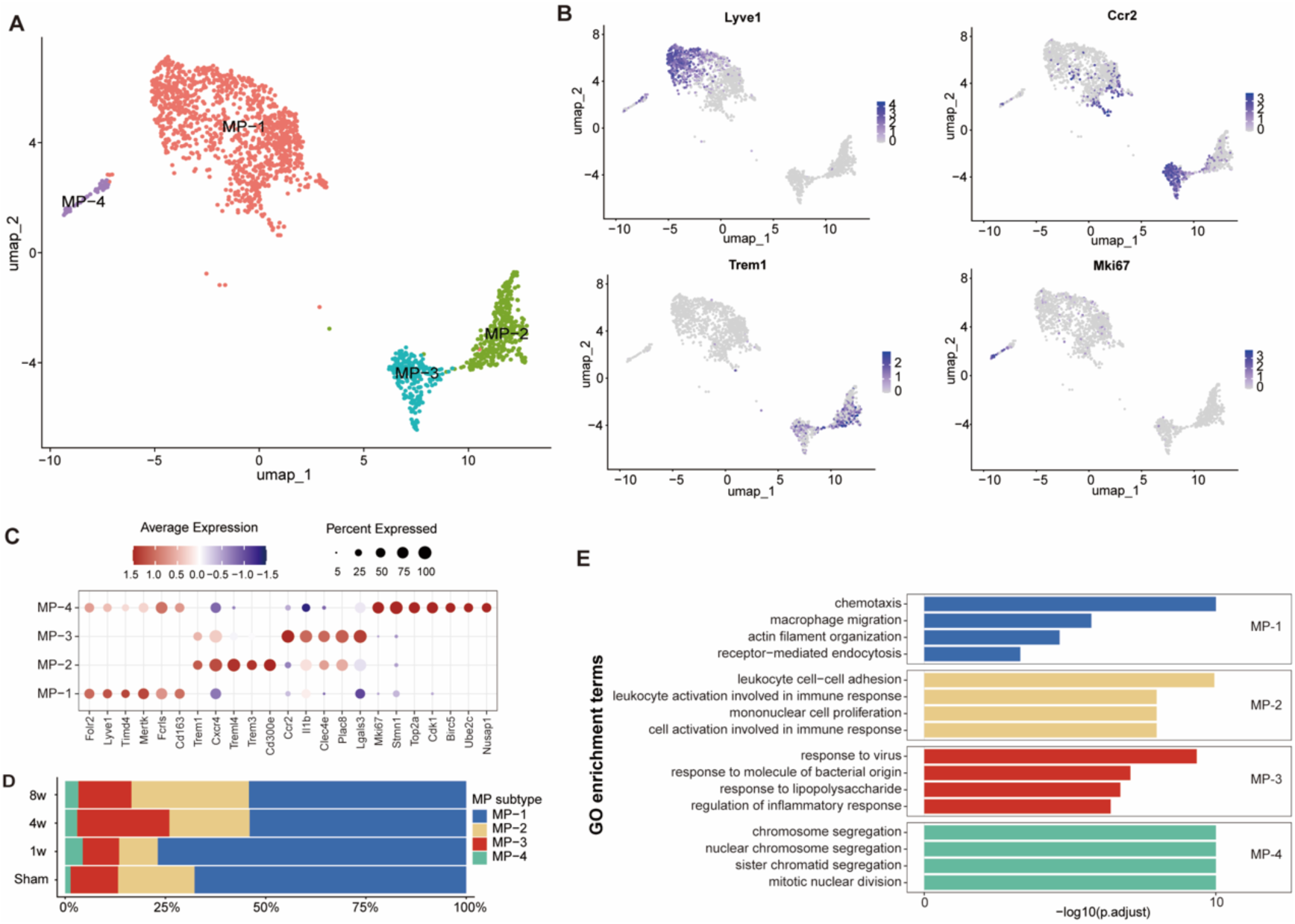
Macrophage heterogeneity and inflammatory remodeling during HFpEF progression. **(A)** UMAP visualization of reclustered macrophages, identifying four macrophage subtypes (MP-1 through MP-4). **(B)** Feature plots showing expression of representative macrophage-state markers, including *Lyve1*, *Ccr2*, *Trem1*, and *Mki67*. **(C)** Dot plot showing marker genes associated with macrophage subtypes. Dot size represents the percentage of cells expressing each marker, and color represents scaled average expression. **(D)** Proportion plot showing relative abundance of macrophage subtypes across control and HFpEF stages. **(E)** GO enrichment analysis showing subtype-specific macrophage programs, including chemotaxis and macrophage migration in MP-1, leukocyte activation and immune-response programs in MP-2, inflammatory and pattern-recognition response pathways in MP-3, and cell-cycle and mitotic programs in MP-4. Bar length represents −log10(adjusted P-value).

Cell composition analysis showed that macrophage states shifted dynamically during HFpEF progression. Compared with control and early HFpEF-1w, the proportion of Lyve1+ MP-1 was reduced at 4w and 8w, whereas inflammatory MP-2 and MP-3 populations increased during HFpEF progression (**Fig. 6D**). MP-4 remained a smaller cycling macrophage subset marked by cell-cycle gene expression throughout the time course. These changes indicate that HFpEF progression is associated with a shift from resident-like Lyve1+ macrophage states toward inflammatory macrophage states, accompanied by a smaller proliferative macrophage compartment.

Functional enrichment analysis further demonstrated that these macrophage subpopulations were associated with distinct biological programs (**Fig. 6E**). MP-1 was associated with migratory and homeostatic functions, including chemotaxis, macrophage migration, actin filament organization, and receptor-mediated endocytosis. MP-2 exhibited enrichment of immune activation-related pathways, including leukocyte cell-cell adhesion, leukocyte activation, mononuclear cell proliferation, and cell activation involved in immune response. MP-3 displayed innate inflammatory and pattern-recognition response signatures, including response to virus, response to bacterial-origin molecules, response to lipopolysaccharide, and regulation of inflammatory response. MP-4 was dominated by cell-cycle-associated pathways, including chromosome segregation and mitotic nuclear division, consistent with its proliferative identity. Together, these findings indicate that HFpEF progression is accompanied by coordinated macrophage remodeling, characterized by reduced representation of resident-like macrophages and expansion of inflammatory states. Their overlap with endothelial and FB immune-response programs suggests a coordinated vascular–immune–stromal remodeling network. This observation prompts us to analyze predicted intercellular communication across the disease time course.

### Intercellular communication analysis predicts stage-dependent EC-immune and macrophage-FB crosstalk

It is well established that extensive intercellular communication exists within the mouse heart^16^. Increasing evidence further suggests that cell-cell communication is markedly enhanced under heart failure conditions^17–19^. Because endothelial, FB, and macrophage analyses revealed coordinated inflammatory, adhesion, and matrix-remodeling programs during HFpEF progression, we next asked whether these cellular changes were accompanied by altered intercellular communication. We performed CellChat analysis to infer ligand-receptor interactions among major cardiac non-CM populations across disease stages. Because CellChat infers potential communication based on co-expression of ligands and receptors, these results are interpreted as predicted signaling interactions rather than as direct evidence of physical or functional cell-cell contact.

CellChat predicted a stage-dependent remodeling of the cardiac intercellular communication network during HFpEF progression. The total number of inferred interactions increased from 2,631 in Ctrl/0w hearts to 3,182 at 1w, declined at 4w, and increased again at 8w. Overall inferred interaction strength followed a similar pattern, increased from 80.9 in Ctrl/0w hearts to 113.4 at 1w, decreasing at 4w, and increasing again (105.2) at 8w, indicating enhanced predicted communication activity during both early and advanced disease stages (**Fig. 7A**). Cell-type-level analysis further suggested that ECs and FBs were major contributors to the increased inferred interaction strength, with additional contributions from SMCs, MPs, NEUTs, NK cells, and PLTs (**Fig. 7B**). These findings suggest that HFpEF progression is accompanied by dynamic remodeling of the cardiac non-CM communication network, with vascular and stromal compartments acting as prominent signaling nodes.

**Figure 7.**
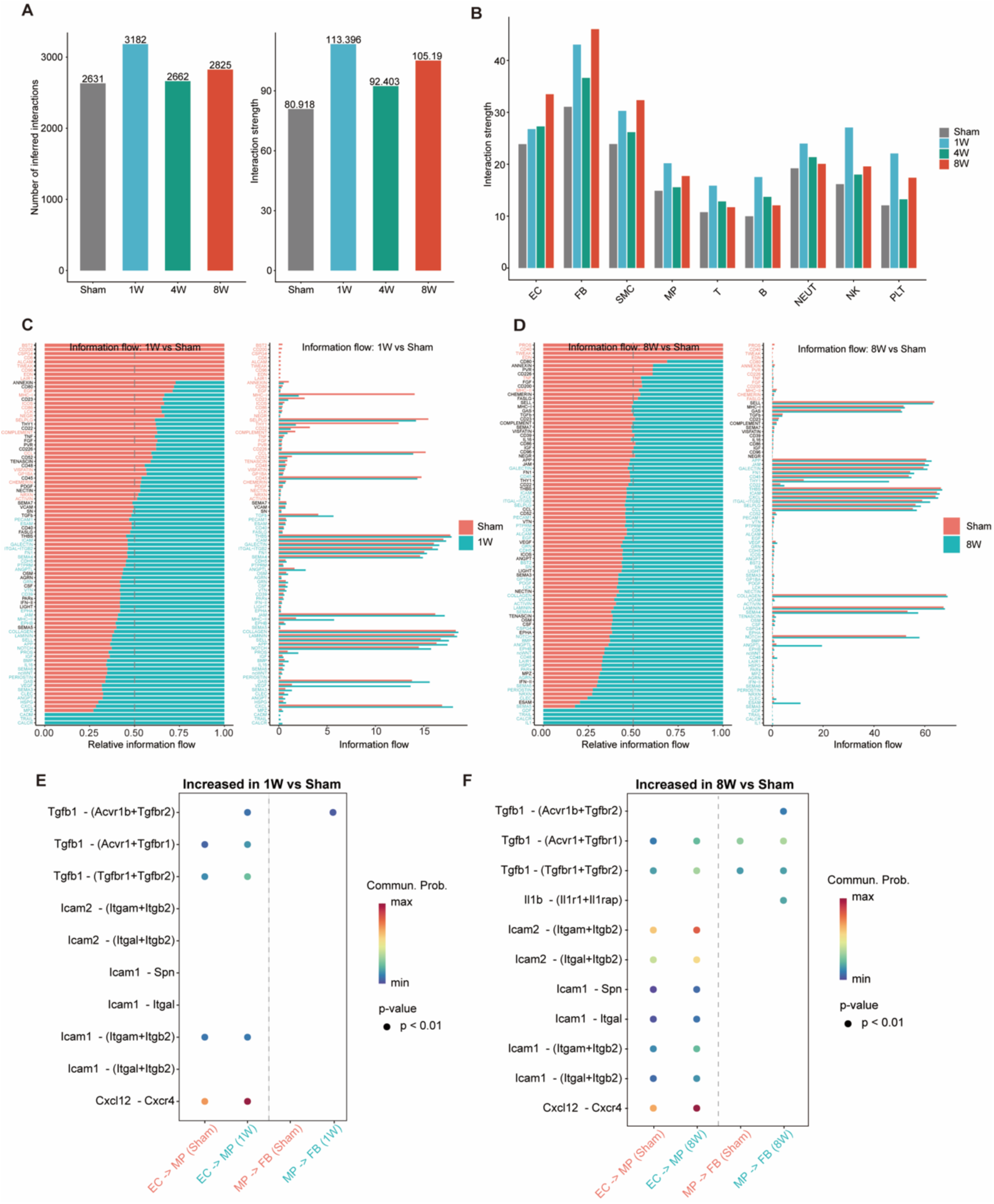
Intercellular communication networks during HFpEF progression. **(A)** Total number of inferred ligand-receptor interactions and overall inferred interaction strength across control and HFpEF stages. **(B)** Inferred interaction strength by major cell population across disease stages, showing dynamic predicted communication involving endothelial, fibroblast, smooth muscle, macrophage, lymphoid, neutrophil, NK cell, and platelet populations. **(C)** Relative and absolute information flow of signaling pathways in HFpEF-1W versus control. Information flow was defined as the summed communication probability across all interacting cell-subpopulation pairs. **(D)** Relative and absolute information flow of signaling pathways in HFpEF-8W versus control. **(E)** Selected ligand-receptor pairs predicted to increase at 1 week, highlighting early endothelial-to-macrophage interactions including chemokine and adhesion-related signaling such as CXCL12-CXCR4 and ICAM1-integrin pairs. **(F)** Selected ligand-receptor pairs predicted to increase at 8 weeks, highlighting persistent endothelial-macrophage signaling and later macrophage-fibroblast inflammatory-fibrotic communication, including IL1 and TGF-family interactions.

Network-level analysis further supported stage-dependent remodeling of predicted intercellular communication. At 1w, the differential interaction network and heatmaps showed a broad increase in inferred communication across vascular, stromal, and immune populations, including ECs, FBs, SMCs, macrophages/mononuclear phagocytes, and lymphoid populations, consistent with early global activation of the cardiac non-CM communication network (**Supplementary Fig. 5A and B**). By 8w, increased interactions became more concentrated among vascular and stromal populations, particularly ECs, FBs, and SMCs, while macrophage/mononuclear phagocyte-associated interactions remained evident (**Supplementary Fig. 5C and D**). Together, these network-level changes suggest a shift from broad early vascular– immune activation toward a more organized vascular–stromal and immune–stromal communication program during established disease.

To identify candidate signaling pathways underlying these network-level changes, we next examined pathway-level information flow. At 1w, HFD+L-NAME hearts showed increased predicted signaling pathways associated with immune regulation, adhesion, vascular remodeling, and early matrix responses, including MHC-II, CD40, CD39, GAS/PROS, VEGF, TGFβ, COLLAGEN, LAMININ, FN1, THBS, and POSTN signaling (**Fig. 7C**). At the established 8w stage, inflammatory and adhesion-related predicted pathways became more prominent, including IL1, CCL, CSF, IFN-II, VCAM, ICAM, and JAM signaling. In parallel, pathways associated with fibroblast activation and extracellular matrix remodeling, including PDGF, ACTIVIN, TENASCIN, POSTN, SPP1, COLLAGEN, and TGFβ-related signaling, were increased (**Fig. 7D**). These pathway-level dynamic support a temporal model in which early endothelial-immune and matrix-associated signaling evolves toward later inflammatory-fibrotic signaling.

We then focused on inferred ligand-receptor interactions among ECs, MPs, and FBs, the three compartments highlighted by the preceding analyses. At 1w, enhanced predicted EC-to-MP interactions included chemokine and adhesion-related ligand-receptor pairs such as Cxcl12-Cxcr4, Icam1-Itgam/Itgb2, Icam1-Itgal/Itgb2, Icam1-Spn, and Icam2-Itgam/Itgb2 (**Fig. 7E**). These interactions are consistent with the endothelial inflammatory and leukocyte-adhesion programs identified by scRNA-seq and suggest a potential role of activated ECs in immune-cell recruitment during early HFpEF. At 8w, predicted EC-MP interactions persisted, while MP-to-FB signaling became more evident, with inferred Il1b-Il1r1/Il1rap and predicted Tgfb1 receptor-complex interactions increased, suggesting enhanced macrophage-FB inflammatory and profibrotic communication at later disease stages (**Fig. 7F**).

Based on these computational predictions, we propose a model in which low-grade inflammatory stress promotes endothelial activation during the early stage of HFpEF progression. Activated ECs may upregulate adhesion molecules, including VCAM1, ICAM1, and E-selectin, thereby facilitating immune-cell recruitment into the cardiac microenvironment^20,21^. Recruited or locally expanded macrophages may then contribute inflammatory and profibrotic mediators, including IL-1β and TGF-β, that promote FB activation and matrix remodeling^22,23^. Because these interactions are inferred from ligand–receptor co-expression, they should be interpreted as candidate signaling circuits rather than direct evidence of physical or functional cell–cell communication.

Together, these analyses predict that HFpEF progression is accompanied by stage-dependent remodeling of the cardiac intercellular communication network. Early disease is characterized by increased predicted endothelial–immune communication involving chemokine and adhesion signaling, whereas later disease is marked by increased macrophage–FB and profibrotic signaling pathways.

## Discussion

This study establishes a time-resolved single-cell atlas of cardiac non-CM remodeling during HFpEF progression. By integrating longitudinal phenotyping, whole-heart bulk RNA-seq, and scRNA-seq of cardiac non-CMs, we aligned systemic and cardiac disease progression with cell-type-specific remodeling of the vascular, immune, and stromal compartments. A major finding of this study is that ECs are an early, prominent, and active component of the HFpEF non-CM response, rather than passive responders to systemic stress. In the HFD+L-NAME model, ECs represented the largest recovered non-CM population and displayed marked transcriptional remodeling. Reclustering revealed multiple endothelial states, including capillary, arterial, stress-response, interferon-stimulated, inflammatory, migratory, ECM-remodeling, and proliferative endothelial populations. Disease-upregulated endothelial genes were heavily enriched for inflammatory response, cytokine signaling, interferon-response programs, leukocyte adhesion, chemotaxis, endothelial migration, angiogenesis, and vascular remodeling, supporting the concept that ECs act as active regulators of the cardiac inflammatory and microvascular microenvironment. Exposing human ECs (HUVECs) to HFpEF-mimic stress similarly induced activation and upregulated adhesion genes (ICAM1, VCAM1, CCL2, and CXCL10). Because HUVECs are macrovascular cells of venous origin rather than cardiac microvascular ECs, this reductionist assay does not fully recapitulate the cardiac microvascular environment, but it provides targeted functional support for the single-cell prediction that HFpEF-associated stress can promote an immune-interacting endothelial state. This inflammatory and adhesion-associated endothelial program was also detected in an independent, L-NAME-free HFD+mTAC model, suggesting that these changes are not solely attributable to nitric oxide synthase inhibition and instead appear conserved across mechanistically distinct HFpEF models.

Following this early endothelial activation, FB and macrophage remodeling emerge as distinct, coordinated components of disease progression. While ECs show early immune-interacting programs, FBs exhibit a matrix-remodeling-dominant response, including ECM organization, collagen biosynthesis, and integrin signaling that becomes prominent at later, established disease stages. Subclustering revealed distinct FB heterogeneity, including ECM/angiogenesis-related (FB-1), chemotaxis/mesenchymal (FB-2), and Wnt/muscle-related (FB-3) pathways, suggesting these cells support both fibrotic matrix deposition and immune-stromal remodeling. Macrophage remodeling provides a potential link between early EC activation and this later FB activity. Reclustering identified a transition during HFpEF progression where homeostatic, resident-like Lyve1/Folr2/Timd4/Mertk-expressing macrophages decline, while inflammatory states (CCR2/TREM1 and TREM-associated) specialized in leukocyte activation expand. Consistent with this cellular shift, ligand–receptor analysis predicted stage-dependent communication circuits: early disease is associated with increased predicted EC–macrophage interactions (ICAM- and CXCL12-associated signaling), transitioning to more prominent predicted macrophage–FB crosstalk (IL1- and TGFβ-family interactions) at later disease stages. Because ligand–receptor analysis is computational, it does not establish protein-level signaling, spatial proximity, or functional interaction, and these circuits should be interpreted as candidates for future mechanistic investigation rather than demonstrated signaling events. This temporal sequence was translationally supported by human HFpEF snRNA-seq data, where human ECs and FBs exhibited parallel inflammatory, vascular activation, and matrix-remodeling pathways. Because this human dataset is cross-sectional, it should be interpreted as supportive translational evidence rather than direct validation of the temporal sequence observed in mice.

Despite these insights, several technical and experimental limitations must be considered. Although the HFD+L-NAME and HFD+mTAC models capture key features of HFpEF, they cannot fully reproduce the clinical heterogeneity of human disease, and cross-species comparisons remain influenced by baseline differences in sample sources, comorbidities, and sequencing platforms. Because this study focused exclusively on viable non-CMs, cardiomyocyte (CM) transcriptional remodeling was not captured within the same single-cell framework, necessitating future work that integrates spatial omics or CM-specific transcriptomics. Methodologically, this study used only male mice; future validation in female cohorts is needed to evaluate sex-based differences. For scRNA-seq, non-CMs from 3 hearts were pooled into a single sample at each time point, precluding statistical modeling at the ‘pseudobulk’ level across biological replicates. Consequently, differential expression results derived from cell-level analyses should be viewed as descriptive findings that primarily serve to generate hypotheses. To mitigate this limitation, we cross-validated key features of EC activation using cross-model comparisons, human snRNA-seq data, tissue-level validation, and *in vitro* experiments incorporating independent biological replicates.

Single-cell dissociation and capture efficiency can also influence apparent cell proportions. Therefore, variations in cell abundance are interpreted as changes in the recovered viable cell population rather than direct measurements of in vivo abundance. Although predominance of ECs among recovered non-CMs is biologically plausible and supported by previous cardiac composition studies^16,24^, dissociation-associated scRNA-seq can introduce compositional bias^16^. Enzymatic digestion, tissue processing methods, cardiomyocyte depletion, filtration, cellular fragility, and droplet-based capture can influence apparent cell proportions. The direction of this bias varies across studies: some protocols may favor endothelial recovery^25,26^, while other studies have found ECs to be relatively sensitive to dissociation-induced stress, potentially leading to their underrepresentation (e.g., lower proportions) rather than enrichment in the sample^27^. Therefore, we interpret predominance of ECs as a characteristic of the cell recovery process rather than a direct measure of absolute abundance in vivo. Importantly, our conclusions regarding endothelial remodeling are supported not only by the proportion of recovered ECs but also by evidence from transcriptional activation, subpopulation remodeling, cross-model and human comparisons, tissue validation, and in vitro functional assays. Furthermore, intercellular communication inferred via CellChat does not establish protein-level signaling or functional causality; cultured human ECs cannot fully recapitulate the properties, tissue mechanical environment, or multicellular interactions of cardiac microvessels in vivo; and cross-species comparisons may be influenced by differences between human and mouse datasets regarding sample sources, disease stages, comorbidities, drug exposures, and sequencing platforms.

In summary, this study defines a time-resolved framework for cardiac non-CM remodeling during HFpEF progression. Our findings highlight early endothelial activation as a central component of a coordinated vascular–immune–stromal remodeling program that includes inflammatory macrophage remodeling, FB matrix activation, and stage-dependent intercellular communication. This atlas provides a foundational resource for future mechanistic studies to define how these cellular states and intercellular communication networks drive the structural and functional progression of HFpEF.

## Data availability

The bulk RNA-seq and scRNA-seq datasets generated in this study will be deposited in the Gene Expression Omnibus (GEO) before completion of peer review. In the interim, data are available from the corresponding author upon reasonable request.

## Acknowledgements

We thank the Genomics, Epigenomics and Sequencing Core (GES Core) at the University of Cincinnati.

## Methods

### Animals

Male wild-type mice (C57BL/6J, 8 weeks of age) were purchased from the Jackson Laboratory (Cat. No. 000664). Mice were housed under standard conditions with free access to food and water. All animal procedures were approved by the University of Cincinnati Institutional Animal Care and Use Committee (IACUC) and were performed in accordance with institutional guidelines. Chemical and biosafety-related procedures were reviewed and approved by the institutional biosafety committee.

### Mouse models

HFpEF murine models were induced using two multi-hit mouse models: metabolic stress (high-fat diet, HFD) plus chemical (Nω-nitro-L-arginine methyl ester hydrochloride, L-NAME) (Sigma-Aldrich, Cat. No. N5751-25G) and HFD plus mechanical stress (mild transverse aortic constriction, mTAC).

For the HFD+L-NAME HFpEF model, mice were fed a 60 kcal% HFD (Cat. No. D12492i) supplemented with L-NAME (1,500 mg/kg diet, Cat. No. D21092202i). Mice were maintained on the HFD+L-NAME diet for up to 12 weeks. Control mice were maintained on normal chow (referred to as Ctrl/0 week). Mice were analyzed at Ctrl/ 0 week (0w), 1 week (1w), 4 weeks (4w), 8 weeks (8w), and 12 weeks (12w), depending on the experimental endpoint.

For the HFD+mTAC HFpEF model, mice (8 weeks of age) underwent mTAC and were maintained on 60 kcal% HFD (Cat. No. D12492i) up to 8 weeks. Control mice with sham surgery were maintained on normal chow (referred to as Sham). Mice were analyzed at Sham, 1w, 4w, and 8w, depending on the experiment endpoint.

### mTAC surgery

A 25-gauge needle was used to induce HFpEF as previously described^28–30^. Mice were anesthetized using inhaled isoflurane, using 2–3% for induction and 1-2% for maintenance. After being mechanically ventilated, the chest was opened and the transverse aorta was exposed. A 6-0 silk suture was passed beneath the transverse aorta, and a 25-gauge needle was placed parallel to the vessel. The suture was tied securely around both the aorta and the needle, after which the needle was promptly withdrawn to leave a fixed stenotic lumen diameter. After closing the chest cavity, the mice were allowed to recover under appropriate postoperative monitoring.

### Intraperitoneal glucose tolerance test

For intraperitoneal glucose tolerance testing, mice were fasted for 4 hours before glucose administration^31^. D-glucose (Sigma-Aldrich, Cat. No. G8270) was injected intraperitoneally at a dose of 1.5 g/kg body weight. Blood glucose was measured from tail-vein blood at baseline (0 minutes) before glucose injection and at 15, 30, 60, 90, and 120 minutes after glucose injection. Glucose levels were measured using an AimStrip Plus Blood Glucose Meter with AimStrip Plus Blood Glucose Test Strips (Germaine Laboratories, Cat. Nos. 37321 and 37350).

### Exercise tolerance testing

Exercise capacity was assessed using an AccuPacer treadmill system (Omnitech Electronics, Inc.). Mice were acclimated to the treadmill for 2 consecutive days before testing. On day 1, mice were placed on the treadmill for 5 minutes without running, followed by 5 minutes at 2.5 m/min. On day 2, mice ran for 5 minutes at 2.5 m/min, followed by 5 minutes at 5 m/min. Then, testing and data collection were performed on day 3 with the treadmill set at a 5-degree incline, starting at 5 m/min for 2 minutes, then 8 m/min for 2 minutes, after which the speed was increased by 2 m/min every 2 minutes until exhaustion. Exhaustion was defined as the mouse remaining on the shock grid for more than 5 seconds despite encouragement. Maximal running time and distance were recorded for analysis.

### Non-invasive blood pressure measurement

Blood pressure was measured using the CODA non-invasive blood pressure system (Kent Scientific Corporation, Torrington, CT). Mice were placed into restraint holders on a warming platform maintained at 35°C and covered with a warming blanket to promote tail blood flow. Tail cuffs were placed and secured according to the manufacturer’s instructions. For each session, mice first underwent 10 acclimatization cycles, followed by 10 measurement cycles. The five highest-quality readings were used for analysis.

### Transthoracic echocardiography

Transthoracic echocardiography was performed using a Vevo 2100 Imaging System equipped with a 40-MHz probe (VisualSonics). Mice were lightly anesthetized with inhaled isoflurane and placed on a heated platform to maintain body temperature. Cardiac images were acquired in two-dimensional long-axis and short-axis views at the level of the maximal left ventricular diameter. Systolic function was assessed by standard measurements of left ventricular dimensions and ejection fraction. Doppler measurements were obtained to evaluate diastolic filling parameters. Image acquisition and analysis were performed as previously described.

### Cardiac Non-cardiomyocyte Single-Cell Isolation

Non-cardiomyocytes were isolated from mouse hearts for single-cell RNA sequencing by enzymatic digestion. Mice were euthanized, and hearts were excised, perfused with PBS, and the atria were removed. Ventricular tissue was minced into a fine slurry and digested in 3 mL of PBS containing Collagenase Type II (1 mg/mL; Worthington, Cat. No. LS004176) and Dispase (1 mg/mL; Gibco, Cat. No. 17105-041). Tissue was incubated at 37°C for around 30 minutes with gentle agitation. Mechanical trituration was performed every 10 minutes using a wide-opening pipette to facilitate gentle dissociation. The resulting cell suspension was filtered twice through a 40-µm cell strainer, washed with PBS, and pelleted by centrifugation. Cells were then resuspended in 0.2% bovine serum albumin (BSA) (Sigma-Aldrich, Cat. No. A9418) in PBS.

### Human endothelial cell culture

Human endothelial cells (hECs, a HUVEC-derived hybrid endothelial cell line; ATCC, Cat. No. CRL-2922) were maintained at 37°C in a humidified incubator with 5% CO₂ according to the manufacturer’s protocol. The cells were cultured in Dulbecco’s Modified Eagle’s Medium (DMEM, ATCC, Cat. No. 30-2002) supplemented with 10% fetal bovine serum (FBS) (ATCC, Cat. No. 30-2020). To mimic an HFpEF in vitro environment, cells were cultured with palmitate (100μM) (Glpbio, Cat. No. GC46103-5), Angiotensin II (1μM) (Glpbio, Cat. No. GP10023-5) for 24 hours, followed by the addition of TNF-α (10ng/ml) (Bio-techne, Cat. No. 210-TA) and IFN-γ (20ng/ml) (Bio-techne, Cat. No. 285-IF) for the final 4 hours. A 0.5% BSA (Glpbio, Cat. No. GC46102) was used as a vehicle control.

### Immune cell–endothelial cell adhesion

The human monocyte THP-1 cells (ATCC, Cat. No. TIB-202) were maintained at 37°C in a humidified incubator with 5% CO₂ according to the manufacturer’s protocol. The THP-1 cells were cultured in suspension in RPMI-1640 medium plus 10% FBS and 0.05 mM 2-mercaptoethanol. To generate GFP-labeled THP-1-derived macrophages, THP-1 cells were cultured in a 6-well plate at a density of 1.2 × 10^6^ cells per well and transduced overnight with adenovirus expressing green fluorescence protein (CMV-GFP) in RPMI-1640 medium (ATCC, Cat. No. 30-2001). The next day, to induce macrophage differentiation, the GFP-containing medium was removed and replaced with complete medium containing phorbol 12-myristate 13-acetate (PMA, 100 ng/mL, Sigma-Aldrich, Cat. No. P8139)^32–34^ for 48 hours. After differentiation, the medium was removed, and cells were gently washed once with PBS. Fresh PMA-free complete medium was added for an additional 24-hour resting period to stabilize the macrophage-like phenotype.

For the adhesion assay, hECs were grown to a confluent monolayer under the indicated experimental conditions. GFP-labeled THP-1-derived macrophages were collected, resuspended in 1.5 mL of HBSS supplemented with Ca²⁺ and Mg²⁺, and added to confluent hEC monolayers at 2.5× 10⁵ cells per well (12-well plate). Co-culture was incubated at 37°C with 5% CO₂ for 15 minutes in 1.5 mL of HBSS supplemented with Ca²⁺ and Mg²⁺. The plate was gently rocked once or twice during incubation to promote even distribution of cells. After incubation, non-adherent cells were removed by gently washing the wells three times with PBS (with Ca²⁺/Mg²⁺). Washing was performed carefully along the wall of the wells to avoid detaching adherent cells. Adherent GFP-positive THP-1-derived macrophages were imaged by fluorescence microscopy, and adhesion was quantified as the number of GFP-positive cells per field from randomly selected fields per well.

### RNA extraction and RT-qPCR

Total RNA of the cells was isolated using Invitrogen™ PureLink™ RNA Mini Kit according to the manufacturer’s protocol (ThermoFisher, Cat. No. 12183018A). Complementary DNA was synthesized using iScript™ Advanced cDNA Synthesis kit (Bio-Rad, Cat. No. 1725038), according to the manufacturer’s instructions. qPCR was performed on a CFX96 Real-Time PCR system (Bio-Rad) using the iTaq Universal SYBR Green Supermix (Bio-Rad, Cat. No. 1725121). The fold changes of each target mRNA expression relative to 18sRNA under experimental and control conditions were calculated based on the threshold cycle (CT) as *r* = 2−Δ(ΔCT), where ΔCT = CT(target)−CT(18S) and Δ(ΔCT) = ΔCT (experimental) − ΔCT(control). The primers for qPCR are listed below,

HomoCXCL10-F: GTGGATGTTCTGACCCTGCT;
HomoCXCL10-R: GGAGGATGGCAGTGGAAGTC;
HomoCCL2-F: CAGCCACCTTCATTCCCCAA
HomoCCL2-R: GGACACTTGCTGCTGGTGAT
HomoICAM1-F: TGACCGTGAATGTGCTCTCC
HomoICAM1-R: TCCCTTTTTGGGCCTGTTGT
HomoVCAM1-F: GGGAGCACTGGGTTGACTTT
HomoVCAM1-R: CATCTCCGTACCATGCCAGT
Homo18srRNA-F: CGGCTACCACATCCAAGGAA
Homo18srRNA-R: GCTGGAATTACCGCGGCT

### Single-cell RNA-sequencing (scRNA-)seq

Two mouse models of HFpEF were employed in this study, including the HFD+L-NAME HFpEF model and HFD+mTAC model. Non-CMs were isolated from mouse hearts and subjected to single-cell RNA sequencing (scRNA-seq). Chromium GEM-X Flex Gene Expression Mouse 16-plex kit (10x Genomics, PN-1000798) was used for GEM generation, reverse transcription of RNA, and cDNA amplification according to the manufacturer’s instructions, and loaded on 10× Genomics Chromium Single Cell Controller to generate barcoded single cells for construction of single-cell cDNA libraries. Targeted cell recovery was 20000 cells. Cell Ranger (v9.0.1) was used for demultiplexing and counting. The 10× Genomics mouse reference genome, refdata-gex-GRCm39-2024-A, was used as reference genome. Data preprocessing and visualization were performed using the R package Seurat (v5.3.0). Cells expressing fewer than 200 genes and genes detected in fewer than three cells were excluded. Doublets were identified using scDblFinder (v1.22.0). The Seurat object was first converted to a SingleCellExperiment object, and doublets were predicted using scDblFinder with default parameters and an expected doublet rate of 5%. Predicted doublets were removed from the dataset, retaining only singlet cells for downstream analyses. In addition, potential doublets were manually inspected based on the co-expression of marker genes from different cell types. After doublet removal, single cells from the control/sham and HFpEF samples were normalized using the NormalizeData function. Highly variable genes were identified using FindVariableFeatures, with the top 3,000 genes selected. The data were scaled using ScaleData, followed by principal component analysis using the selected variable features. To correct for batch effects across different groups, Harmony (v1.2.4) was applied based on the PCA embeddings using the RunHarmony function, with group information specified as the integration variable. The first 30 Harmony dimensions were used to construct the shared nearest neighbor graph using FindNeighbors, and clustering was performed using FindClusters (resolution = 0.5). Uniform Manifold Approximation and Projection (UMAP) was applied using RunUMAP based on the Harmony reduction for visualization.

### Marker gene analysis

Marker genes for each cluster were identified by FindAllMarkers function. Low-quality clusters were removed, and the remaining cells were reclustered. Cell identities were assigned to each cluster based on canonical marker genes using the AddModuleScore function in Seurat, with marker genes curated from the CellMarker 2.0 database and previously published studies. Differential gene expression analysis between Sham, 1, 4 and 8 weeks samples for each cluster were performed using the FindMarkers function in Seurat with the Wilcoxon rank-sum test. Log-normalized counts from the RNA assay were used as input. Genes with a log2 fold change > 0.25, adjusted P value < 0.05, and expression in at least 10% of cells were considered for downstream analysis.

### Subclustering analysis

For subclustering analysis in the HFD+L-NAME HFpEF model, individual cell populations were subsetted from the integrated dataset and reclustered separately. For endothelial cell (EC) subclustering, ECs were subsetted and reclustered using the first 15 principal components with a resolution of 0.45, resulting in 8 distinct EC subtypes. Subtypes were annotated based on previously reported canonical marker genes. For fibroblast (FB) subclustering, FBs were subsetted and reclustered using the first 15 principal components with a resolution of 0.25, resulting in 3 FB subtypes. Macrophage (MP) subclustering was performed similarly using a resolution of 0.05, resulting in 4 MP subtypes. All subtypes were annotated based on canonical marker genes from published studies.

### Cell-type proportion analysis

Cell-type proportions were calculated for each group (Ctrl/0w, 1w, 4w, and 8w) and visualized as stacked bar plots. To summarize differential gene expression across clusters and conditions, the numbers of upregulated and downregulated genes were quantified and visualized using lollipop plots. For GO enrichment analysis, significant genes were selected using a uniform filtering strategy (adjusted P value < 0.05 and log2 fold change > 0.25). For cell subtype characterization, the top-ranked marker genes (top 250 per cluster based on log2 fold change) were used as input for GO enrichment analysis to define subtype-specific functional features and support cell subtype annotation. For disease-associated pathway analysis, upregulated differentially expressed genes identified between disease and control groups were used to assess functional changes associated with disease progression. GO enrichment analysis was performed using the clusterProfiler (v4.17.0) R package. Enriched GO terms were visualized using bar plots and heatmaps, with enrichment significance represented by −log10(adjusted P value), to compare functional differences across cell subtypes, disease conditions, and time points. Dot plot and bar plots representing significantly enriched pathways were generated by ggplot2 (v4.0.3.9) R package. Heatmaps for scRNA-seq data were generated using the DoHeatmap function in Seurat and the ggplot2 R package for advanced visualization. Curated gene signatures representing ECM remodeling, vascular, stromal, Wnt signaling, inflammatory, and interferon-response programs were used to characterize fibroblast subtypes based on mean normalized expression, with cluster-level scores visualized as heatmaps. Mean module scores were visualized as dot plots, with color representing the average module score and dot size indicating the number of cells in each cell type and time point. To assess inflammatory and adhesion-related transcriptional activity across major non-cardiomyocyte populations in both HFpEF models, a gene set comprising genes associated with cell adhesion, chemokine-mediated leukocyte recruitment, inflammatory signaling, interferon responses, and endothelial activation was used to calculate module scores with the Seurat AddModuleScore function. Module scores were calculated at the single-cell level and subsequently averaged for each major cell type at each HFpEF time point (1, 4, and 8 weeks).

### Intercellular communication analysis

Intercellular communication analysis was performed using the CellChat (v1.6.1) R package to infer ligand–receptor (LR) interactions between cell populations, using the curated mouse ligand–receptor interaction database (CellChatDB.mouse) as prior knowledge. CellChat objects were constructed separately for each condition (Sham, 1W, 4W, and 8W), and communication probabilities were computed using the computeCommunProb function with the triMean method. Interactions involving cell groups with fewer than 10 cells were excluded. Signaling pathway-level communication was inferred using computeCommunProbPathway and aggregateNet. CellChat objects were merged to compare the total number of inferred interactions across conditions. Cell type-specific communication activity was quantified by summing incoming and outgoing interaction strength for each cell population. Differential communication between conditions was assessed using netVisual_diffInteraction, while relative information flow across signaling pathways was compared between Sham and individual HFpEF time points using the rankNet function in comparison mode. Pathway-specific and selected LR interactions were further examined using bubble plots and heatmaps, and CellChat-inferred communication probabilities were integrated with differential expression analysis of the corresponding ligand and receptor genes.

### Human snRNA-seq data analysis

Publicly available human cardiac snRNA-sequencing data^10^ were obtained in H5AD format and imported into R using the zellkonverter package. The resulting SingleCellExperiment object was converted into a Seurat object, with CellBender-corrected raw counts retained as the count matrix and the normalized expression matrix used for downstream analysis. Precomputed UMAP coordinates and cell-type annotations provided by the original study were retained for visualization and subsequent disease-associated analyses.

The original cell-type annotations were evaluated using cluster-enriched marker genes identified with the Seurat FindAllMarkers function and established canonical markers. Cell-type proportions in control and HFpEF samples were calculated and visualized using the same approach as described for the mouse scRNA-seq datasets. Differentially expressed genes between HFpEF and control samples were identified within individual cell populations using the Seurat FindMarkers function. Upregulated genes meeting the specified differential-expression thresholds were subjected to GO enrichment analysis using clusterProfiler. Species-specific annotation databases were used for GO analysis, with org.Hs.eg.db applied to human genes and org.Mm.eg.db applied to mouse genes. Enriched human GO terms were visualized using dot plots.

