## Supplemental Figure 1-5 for "Time-Resolved Single-Cell Atlas Reveals Early Endothelial Activation and Stage-Dependent Immune–Stromal Communication in HFpEF"

**Short title:** Time-Resolved Single-Cell Atlas of HFpEF Progression

**Address for Correspondence:**
Wei Huang, MD, PhD
Division of Cardiovascular Health and Disease
Department of Internal Medicine
University of Cincinnati College of Medicine
231 Albert Sabin Way
Cincinnati, Ohio 45267


**Supplementary Figure 1-5**

### Supplementary Figure 1

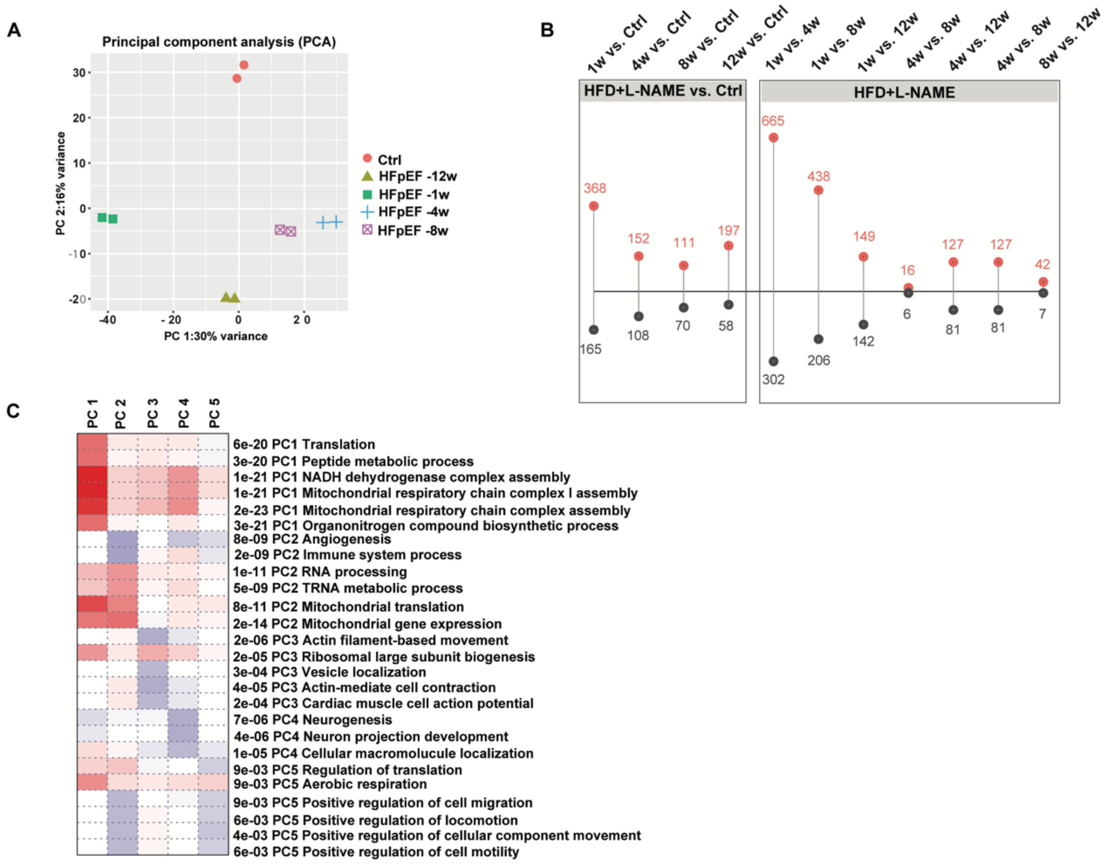


**Supplementary Figure 1. Bulk RNA-seq analysis of heart tissue during HFD+L-NAME HFpEF progression.**

**(A)** Principal component analysis (PCA) of bulk RNA-seq data from control and HFpEF hearts across disease stages.

**(B)** Number of upregulated and downregulated differentially expressed genes (DEGs) across disease-stage comparisons.

**(C)** GO enrichment analysis associated with major principal components, showing pathways related to translation, mitochondrial respiratory chain assembly, angiogenesis, immune system process, RNA processing, mitochondrial gene expression, actin-based movement, and cell motility.

### Supplementary Figure 2

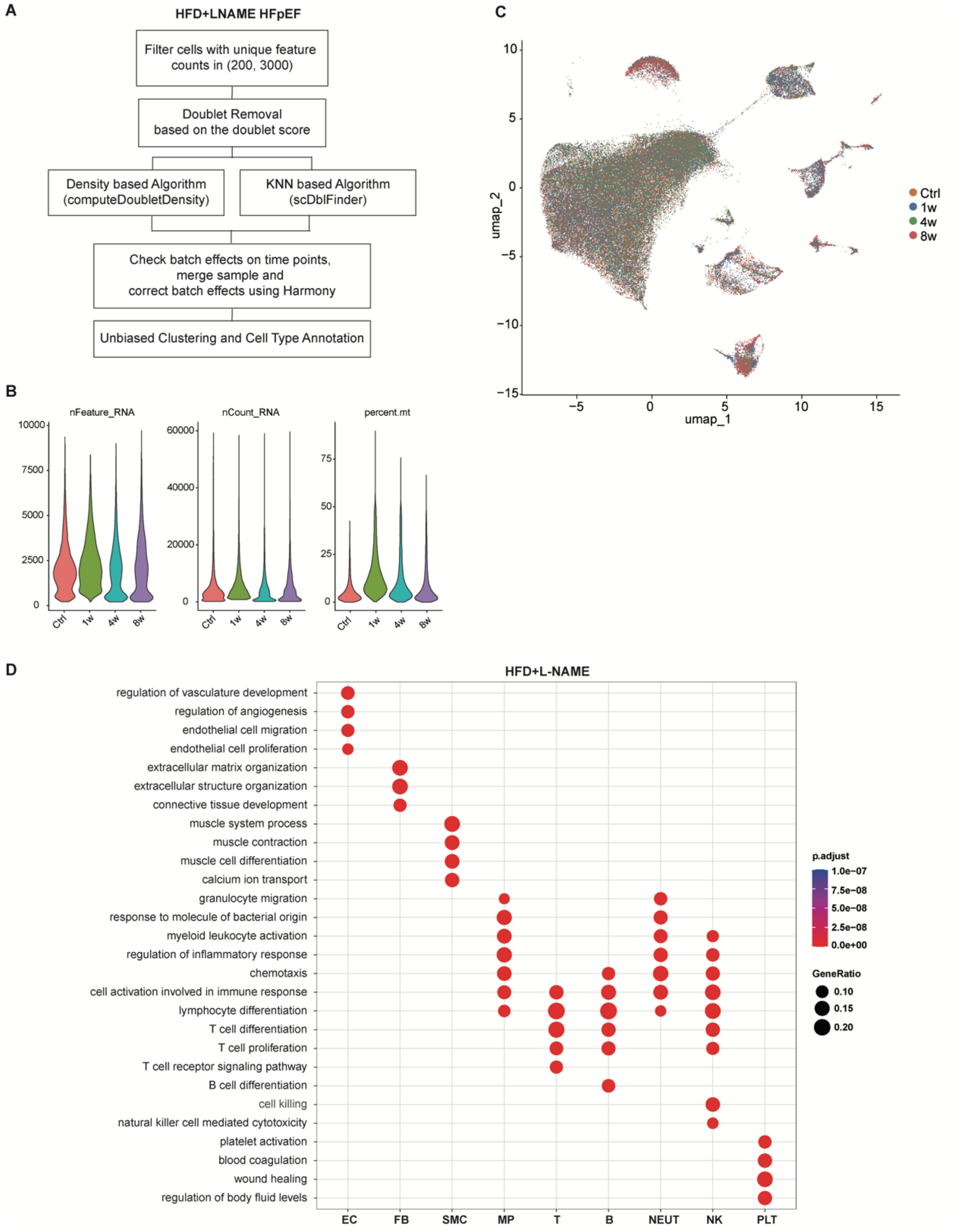


**Supplementary Figure 2. Quality control, batch integration, and cell-type annotation of the HFD+L-NAME scRNA-seq dataset.**

**(A)** Overview of the preprocessing and quality control workflow for scRNA-seq data for HFD+L-NAME cardiac non-CMs, including feature-count filtering (200–3,000 unique features per cell), doublet removal using a consensus density-based (computeDoubletDensity) and KNN-based (scDblFinder) approach, batch-effect assessment and correction across timepoints using Harmony, and unbiased clustering with cell-type annotation.

**(B)** Violin plot showing the distribution of nFeature_RNA (number of unique features (genes) detected in each cell), nCount_RNA (total number of RNA molecules detected per cell), and percent.mt (percentage of mitochondrial gene expression) across different time points after quality-control filtering.
**(C)** UMAP visualization of all recovered non-CMs, grouped by time point, showing no evidence of batch effects and no emergence of distinct clusters across time points.
**(D)** Gene Ontology (GO) enrichment analysis of cell-type-enriched genes across major cardiac non-CM populations (EC, FB, SMC, MP, T, B, NEUT, NK, PLT) in the HFD+L-NAME model. Dot size represents gene ratio, and dot color represents adjusted p-value.

### Supplementary Figure 3

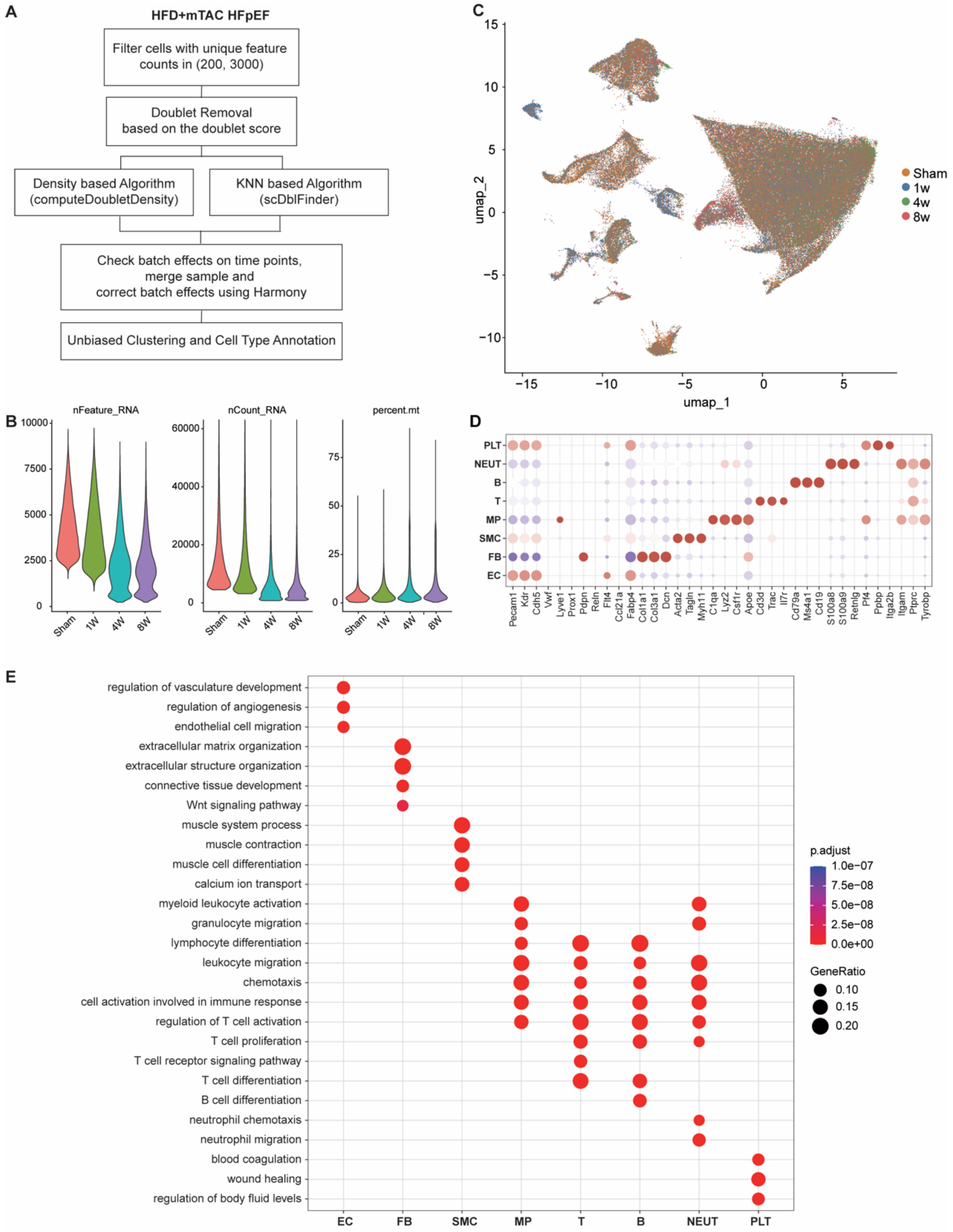


**Supplementary Figure 3. Quality control, batch integration, and cell-type annotation of the HFD+mTAC scRNA-seq dataset.**

**(A)** Overview of the preprocessing and quality control workflow for scRNA-seq data for HFD+mTAC cardiac non-CMs, including feature-count filtering (200–3,000 unique features per cell), doublet removal using a consensus density-based (computeDoubletDensity) and KNN-based (scDblFinder) approach, batch-effect assessment and correction across timepoints using Harmony, and unbiased clustering

with cell-type annotation.
**(B)** Violin plot showing the distribution of nFeature_RNA (number of unique features (genes) detected in each cell), nCount_RNA (total number of RNA molecules detected per cell), and percent.mt (percentage of mitochondrial gene expression) across different time points after quality-control filtering.
**(C)** UMAP visualization of scRNA-seq data, grouped by time point, showing no evidence of batch effects and no emergence of distinct clusters across time points.
**(D)** Dot plot showing canonical marker gene expression used to annotate major cell populations in the HFD+mTAC dataset, including EC, FB, SMC, MP, T cells, B cells, NEUT, and PLT. Dot size indicates the percentage of cells expressing each marker, and color indicates scaled average expression.
**(E)** Gene Ontology (GO) enrichment analysis of marker genes defining each of the eight annotated cell populations. Dot size represents gene ratio, and dot color represents adjusted p-value.

### Supplementary Figure 4

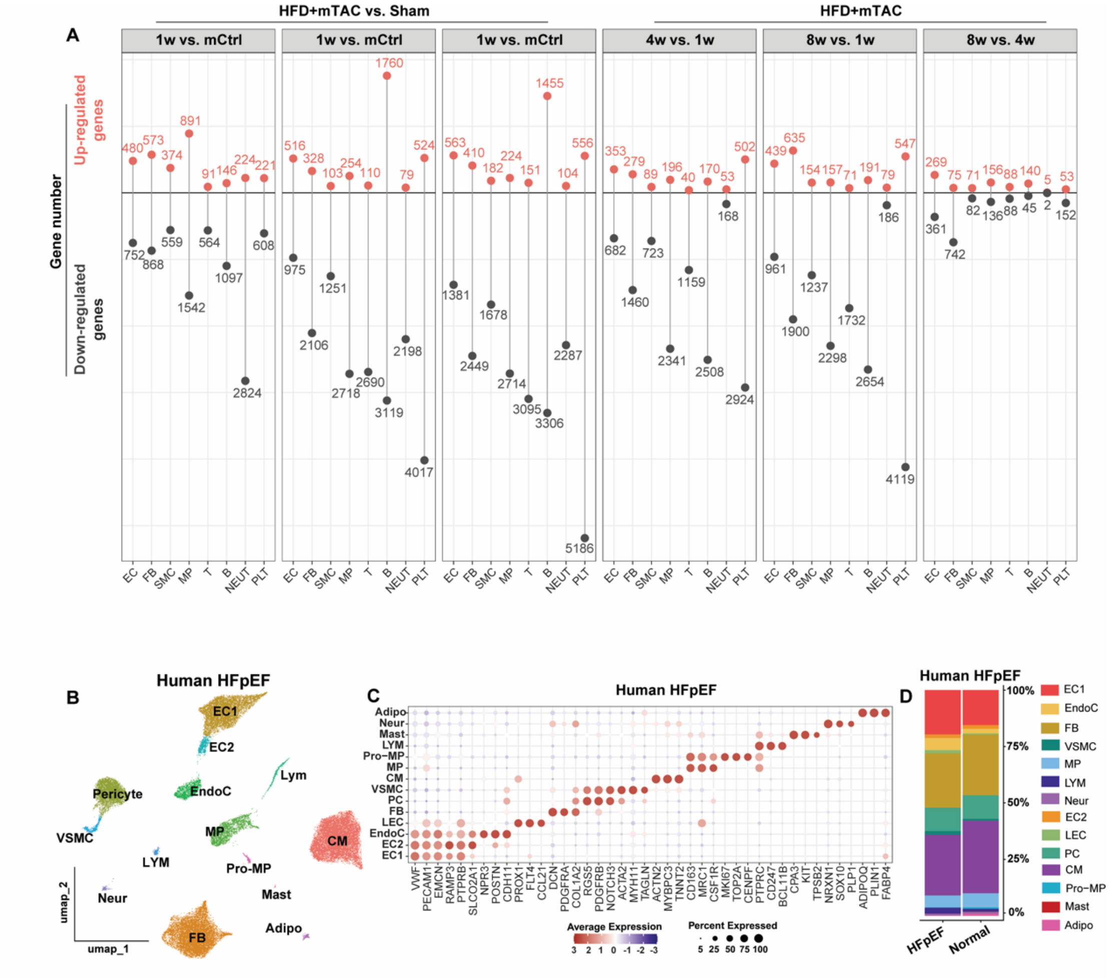

**Supplementary Figure 4. Differential gene expression across HFD+mTAC disease stages and annotation of human HFpEF snRNA-seq data.**

Cross-species support for endothelial and fibroblast remodeling in human HFpEF using single-nucleus RNA-seq (snRNA-seq).

**(A)** Lollipop plot showing upregulated and downregulated DEGs across major cell populations in HFD+mTAC and disease-stage comparisons. DEGs were defined using adjusted P < 0.05 and log fold change > 0.25.

**(B)** UMAP visualization of public human HFpEF single-nucleus RNA-seq data showing major cardiac cell populations, including endothelial cell subsets, fibroblasts, pericytes, vascular smooth muscle cells, cardiomyocytes, macrophages, proliferating macrophages, lymphoid cells, mast cells, neuronal cells, and adipocytes.

**(C)** Dot plot showing canonical marker gene expression used to annotate human cardiac cell populations. Dot size represents the percentage of cells expressing each marker, and color represents scaled average expression.

**(D)** Proportion plot showing relative abundance of recovered human cardiac cell populations in HFpEF and normal/control samples.

### Supplementary Figure 5

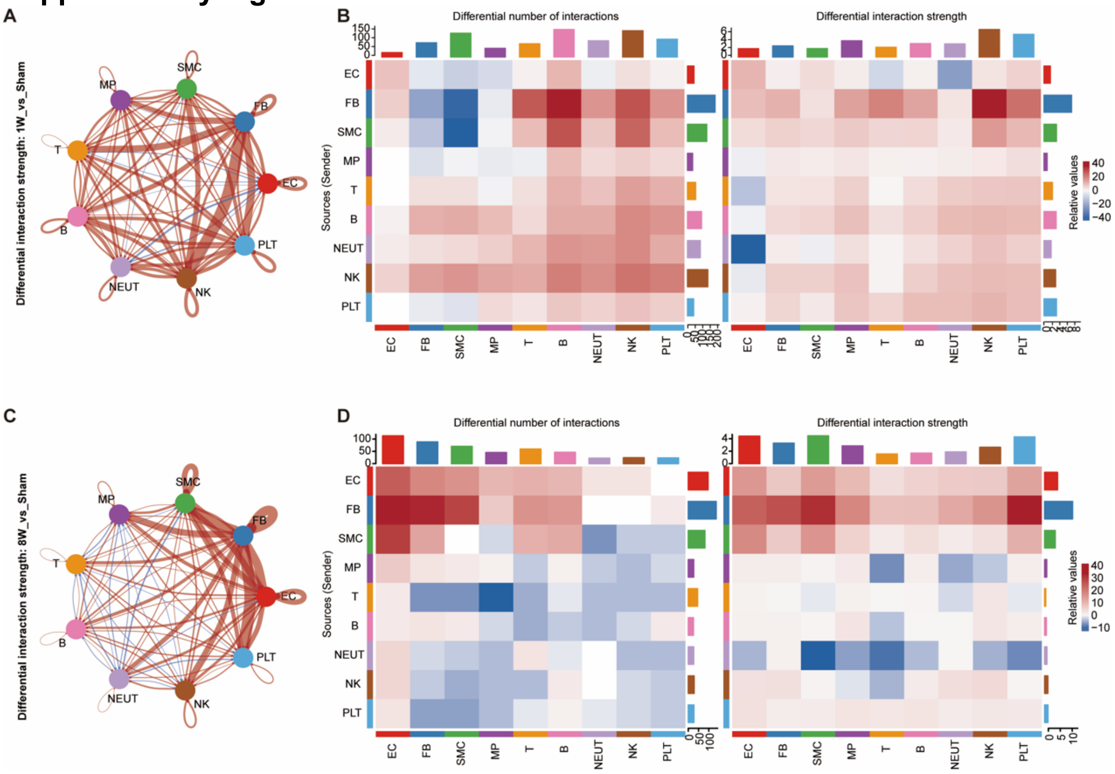


**Supplementary Figure 5. Stage-dependent remodeling of predicted intercellular communication during HFpEF progression.**

**(A)** Differential interaction-strength network comparing 1-week HFpEF hearts with Sham controls. Red edges indicate increased inferred interaction strength at 1 week relative to Sham, whereas blue edges indicate decreased interaction strength. Edge width is proportional to the magnitude of the difference, and node colors denote the indicated cell populations.

**(B)** Heatmaps showing differences in the inferred number of interactions (left) and overall interaction strength (right) between sender cell populations (rows) and receiver cell populations (columns) at 1 week relative to Sham. Red indicates an increase and blue indicates a decrease. Marginal bar plots summarize the overall changes associated with each sender and receiver population.

**(C)** Differential interaction-strength network comparing 8-week HFpEF hearts with Sham controls.

**(D)** Heatmaps showing differences in the inferred number of interactions (left) and overall interaction strength (right) between sender and receiver cell populations at 8 weeks relative to Sham.
